# Population divergence in maternal investment and embryo energy use and allocation reveals adaptive responses to cool climates

**DOI:** 10.1101/2022.12.07.519527

**Authors:** A.K. Pettersen, S. Ruuskanen, Andreas Nord, J. F. Nilsson, M.R. Miñano, L. J. Fitzpatrick, G.M. While, T. Uller

## Abstract

The thermal sensitivity of early life stages can play a fundamental role in constraining species distribution. For egg-laying ectotherms, cool temperatures often extend development time and exacerbate developmental energy cost. Despite these costs, egg laying is still observed at high latitudes and altitudes. How embryos overcome the developmental constraints posed by cool climates is crucial knowledge for explaining the persistence of oviparous species in such environments and for understanding thermal adaptation more broadly. Here, we studied maternal investment, and embryo energy use and allocation in wall lizards spanning altitudinal regions, as potential mechanisms of local adaptation to development in cool climates. Specifically, we compared population-level differences in (1) investment from mothers (egg mass, embryo retention and thyroid yolk hormone concentration), (2) embryo energy expenditure during development, and (3) embryo energy allocation from yolk towards tissue. We found evidence that energy expenditure was greater under cool compared with warm incubation temperatures. Females from relatively cool regions did not compensate for this energetic cost of development by producing larger eggs or increasing thyroid hormone concentration in yolk. Instead, embryos from the high-altitude region used less energy to complete development, i.e., they developed faster without a concomitant increase in metabolic rate, compared with those from the low-altitude region. Embryos from high altitudes also allocated relatively more energy towards tissue production, hatching with lower residual yolk:tissue ratios than low-altitude region embryos. These results suggest that local adaptation to cool climate in wall lizards involves mechanisms that regulate embryonic utilisation of yolk reserves and its allocation towards tissue, rather than shifts in maternal investment of yolk content or composition.

## Introduction

Species distributions are often strongly associated with climate, and recent range expansions suggest that low temperature can be a limiting factor (Calosi et al., 2010; Crozier, 2003; Osland & Feher, 2020). For egg-laying ectotherms such as reptiles, embryonic development is perhaps the most critical life stage since embryos have limited opportunity to behaviourally thermoregulate (Hamdoun & Epel, 2007); but see Du et al., (2011). In such taxa, cold nest temperatures generally extend development times and delay hatching, such that juveniles may start feeding later in the reproductive season, which often is associated with low fitness (Ji & Braña, 1999; While et al., 2015). Nevertheless, egg-laying species have repeatedly, and sometimes rapidly, colonised cooler climates. Identifying the mechanisms by which populations overcome physiological limits is therefore crucial for our understanding of thermal adaptation and their effects on range margins (Shine, 1999).

As a rate-limiting factor of biological systems, incubation temperature affects two processes that determine total energy expenditure during the embryonic stage, or the “cost of development” – the rate of energy expenditure (estimated as metabolic rate; hereafter referred to as *MR*) and the length of the developmental period from fertilisation until hatching (development time; *D*) (Marshall et al., 2020). A decrease in temperature generally decreases *MR* but extends *D*. However, evidence so far suggests that for the majority of ectotherms, cool temperatures extend *D* more than they reduce *MR*, resulting in developmental energy costs that are seemingly greatest at cool temperatures (Pettersen et al., 2019). The implications of this asymmetry between the temperature dependence of *D* and *MR* is that offspring developing at low temperature will deplete yolk reserves and either die before hatching, or hatch with little or no residual energy (DuRant et al., 2011; While et al., 2015). Alternatively, greater cost of development may result in reduced tissue production, and therefore lower mass at hatching (Suarez et al., 1996).

Maternal effects likely play a key role in adaptation to cool climates (Badyaev, 2009; Burgess & Marshall, 2011; Moore et al., 2019; Pettersen et al., 2019). For example, mothers may help to offset the high costs of development in cool climates by either directly increasing energy investment into offspring or reducing the energy embryos require to complete development. First, mothers may increase the size, and therefore energy investment, of their offspring. Experimental work has shown that cooler conditions select for greater per-offspring investment, and that mothers reared under cooler temperatures do indeed produce larger offspring – also known as the offspring-size temperature relationship (Atkinson et al., 2001; Bownds et al., 2010; Yampolsky & Scheiner, 1996). Assuming energy content scales at least isometrically with egg size, this relationship may be an adaptive response to offset costlier development under cooler conditions (Pettersen et al., 2019). Second, mothers may retain their embryos through a longer proportion of development – effectively reducing development time and the cost of development by maintaining embryos at warmer maternal body temperatures (Rodríguez-Díaz & Braña, 2012; Shine, 1989). Third, mothers may alter the composition of the egg yolk that fuels development (Hsu et al., 2019; Valcu et al., 2019). Moreover, past studies on wild bird populations have shown that yolk steroid and thyroid hormones in particular can advance hatching dates, thereby offering potentially promising candidate mechanisms for local adaptation of developmental rate (T. G. G. Groothuis & Schwabl, 2008; Ruuskanen & Hsu, 2018).

There is a growing body of evidence in fish, amphibians, and reptiles that populations in cool climates have evolved faster developmental rates, such that they hatch at comparably similar dates, to populations in warmer climates (known as countergradient variation; (Conover & Present, 1990; Laugen et al., 2003; Pettersen, 2020)). Local adaptation in developmental rates is most easily exemplified by common garden studies, whereby individuals from cold-adapted populations (e.g., high latitudes and altitudes) develop faster than those from warm-adapted (e.g., low latitude and altitude) populations when placed under common conditions (Laugen et al., 2003). Despite clear evidence that countergradient variation represents local adaptation to cool climates (Pettersen, 2020), its underlying proximal drivers are yet to be identified. It is also unclear whether faster developmental rates necessitate increased metabolic rates and therefore, cost of development.

Here we study population divergence in maternal investment, and embryo energy use and allocation across a climatic gradient in the widespread common wall lizard, *Podarcis muralis* (Figure 1A) from the western Italian coast eastward into the Apennine mountains in central Italy. Wall lizards are abundant across this region, which spans >800m in altitude and represent a transition in climatic regime from warm (low altitude) to cool (high altitude) climate (Ruiz Miñano et al., 2022). There is likely no single mechanism acting to compensate for the greater energy cost associated with embryonic development in cool climates, but rather a suite of complementary mechanisms. First, mothers from high altitudes may offset the greater cost of development by investing in larger eggs, retain embryos for a greater proportion of development, effectively accelerating development of their offspring, or alter yolk thyroid hormone concentration to enable faster development. Second, energy use by developing embryos may be reduced by evolving either faster developmental rates, lower metabolic rates, or both. Third, relative energy allocation from yolk reserve towards tissue may increase in embryos from cool climates. Hatching at a larger body size by converting proportionally more yolk towards tissue, could be a viable response in cool climates to aid post-hatching survival (Atkinson, 1996). To test for the presence of these potential adaptations to cool climates, we measure differences for lizards from three climatic ‘regions’ in *i*) maternal investment (egg size, embryo retention, yolk thyroid hormone concentration), and manipulate developmental environments to measure *ii*) thermal dependence of embryo energy use (development time and metabolic rate) and *iii*) embryo energy allocation (tissue to total embryo mass, hatchling mass and length) across cool and warm incubation temperatures.

**Figure 1.**
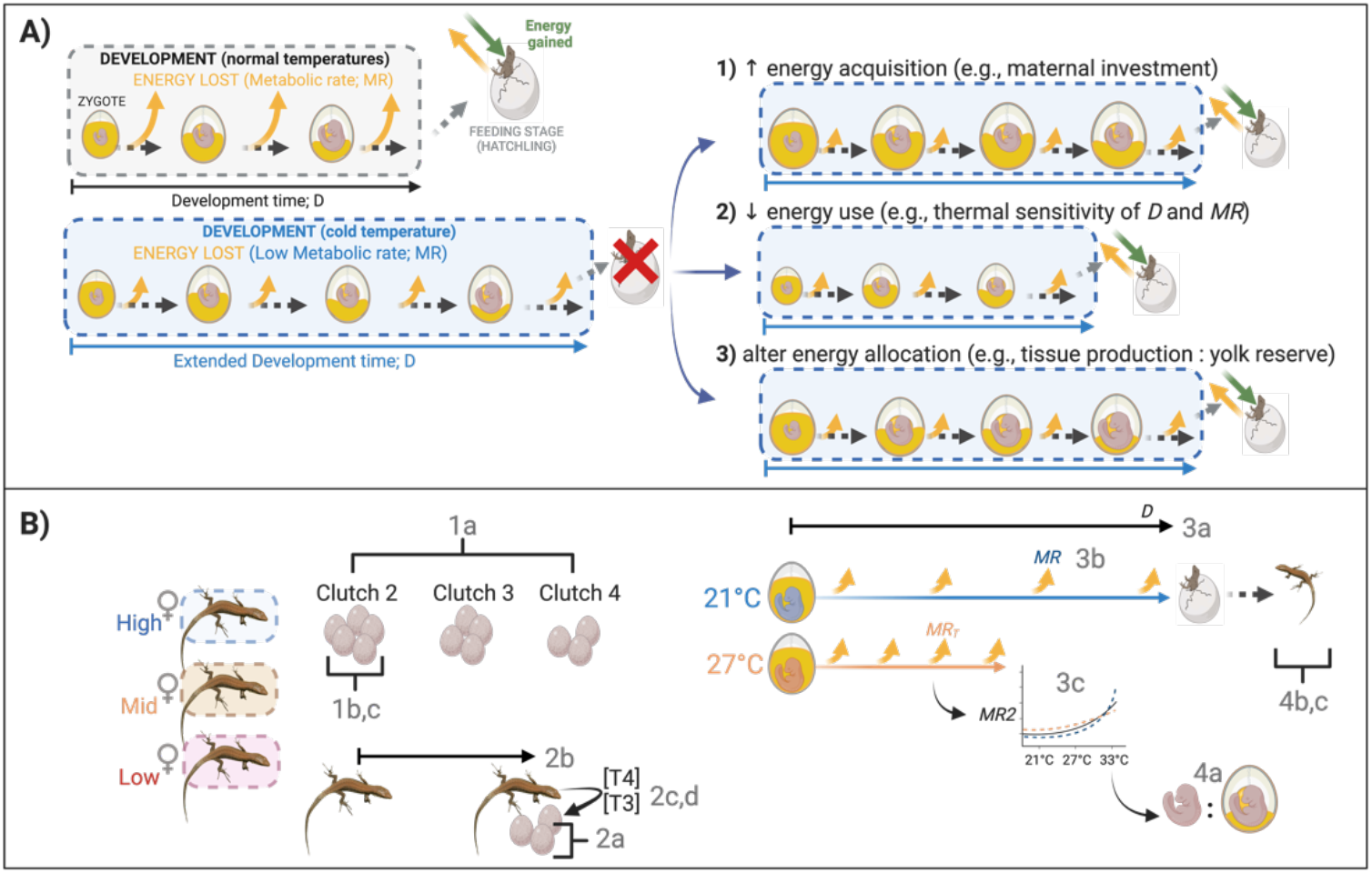
A) Conceptual diagram of predicted acute effects of cool developmental temperatures on the cost of development and potential evolutionary responses, and B) Experimental design. A) Energy use during development (the ‘cost of development’) is a product of metabolic rate and development time and may influence residual yolk reserves at hatching and/or the conversion of yolk reserves into tissue (structure), and therefore survival. Cooler developmental temperatures are often associated with higher costs of development due to a greater thermal sensitivity of development time (resulting in delayed hatching) relative to metabolic rate. Three potential mechanisms by which embryos and mothers can counteract the effects of cool incubation temperature on the costs of development – 1) increase embryo energy acquisition via increased investment from mothers, 2) decrease embryo energy use (and therefore the requirement for increased maternal investment) by reducing development time (*D*), metabolic rate (*MR*), or both, or 3) alter embryo energy allocation to tissue relative to yolk reserve at hatching. B)Experimental split-clutch design for testing the effects of altitude Region (High, Mid, Low), incubation treatment (21 °C or 27 °C), and Clutch order (2^nd^ – 4^th^) on maternal investment (Results section 2a-d), use (3a-3c), and allocation (4a-c).

## Materials and Methods

### Study system

*Podarcis muralis*, the common European wall lizard, is a small (50-70mm snout-to-vent; SVL length), oviparous lacertid. It is widespread across southern Europe and, in contrast to other *Podarcis* lizards, the range extends north of the Mediterranean basin. In the Mediterranean, *P. muralis* has also colonised high-altitude regions, including the Pyrenees, Apennines, and the Alps, occasionally extending >2400m above sea level, where low air, and therefore soil nest temperatures, appear to impose a limit on the species’ distribution (Sindaco et al., 2006). The species has also successfully established at higher latitudes after repeated, independent human introductions of lizards into north-western Europe and north America over the last century (Engelstoft et al., 2020; Michaelides et al., 2013, 2015; Schulte et al., 2012).

### Field collection and animal husbandry

Adult wall lizards (62 females and 38 males) were captured in the field at the start of the reproductive season (March – April 2019) across three regions in central Italy. These locations represent three climatic regimes: low altitude (~130 m; approximate latitude/longitude decimal degrees: 42.070°/ 11.954°), mid altitude (~600 m; 42.153°/12.860°), and high altitude (~1000 m; 42.279°/13.189°); see Table S1 for climatic differences among populations (Lembrechts et al., 2022). Lizards were brought to the animal facilities at Lund University, Sweden, where they were housed in cages (590 × 390 × 415 mm; depth × width × height) with one to two females, and one male, all from the same climatic region. The cages contained sand substrate, a sandbox for egg laying, shelter (bricks and logs), water bowl, basking lamp (60W; 8 h per day), and UV light (EXO-TERRA 10.0 UVB fluorescent tube; 6 h per day). This created a thermal gradient within the cages of 24-40°C while the basking lamps were switched on. Night temperature was set to 20°C and the light cycle to 12 L: 12D during the reproductive season. Lizards were fed mealworms or crickets daily. Sand boxes were inspected daily for clutches laid overnight to ensure egg sampling within 24 h post oviposition. Number of eggs used for experiments per population and female are provided in Table S2.

### Reproductive output measures

Upon egg laying, clutch number (*P. muralis* lay multiple clutches per year; see Results Section 1a) and clutch size (Section 1b), were recorded, and clutch mass (Section 1c) and maternal body mass were measured to the nearest 0.01g (Ohaus Scout SKX Precision Balance). Over the reproductive season (April – August 2019), females laid between 2-5 clutches (62 females, mean: 3.53 ± SD: 1.08) with clutch sizes ranging between 2-10 eggs (62 females, mean: 5.00 ± SD: 1.18). All data on first clutches (except clutch number) was excluded to reduce potential environmental effects from the field confounding our results, and because a small number of females (n = 4 out of 62) had already oviposited in the field. Only 10 females laid a fifth clutch and to maintain a balanced design, data on fifth clutches were also omitted. For clutches 2-4, a split-clutch design was used: one egg per clutch was frozen at −80 °C immediately to determine developmental stage at oviposition and for hormone analysis (see Maternal investment measures below), and the remaining eggs were assigned evenly among the incubation treatments.

### Incubation treatments

Eggs from clutches of known parentage were alternately allocated to one of two incubation temperature treatments: 21 °C (cool) or 27 °C (warm), representing soil temperatures likely to be commonly experienced by eggs in nests of high and low altitudes in Italy, respectively (While et al., (2015), Table S1). Development at constant 21 °C is close to the lowest possible constant temperature of sustained development in *P. muralis* while 27 °C is considered benign (Braña & Ji, 2000; Van Damme et al., 1992; While et al., 2015). Eggs were placed into small plastic containers filled with two-thirds with moist vermiculite (5:1 vermiculite to water volume ratio) and sealed with clingfilm, before being placed into incubators throughout development (except for brief periods during metabolic rate measurements, see details below).

### Maternal investment measures

Per offspring investment from mothers was measured as egg mass (Results Section 2a), extent of embryo retention (stage of embryo development at oviposition; Results Section 2b), and egg yolk thyroid hormone concentration (Results Section 2c, 2d). Egg mass and maternal body mass was measured upon oviposition to the nearest 0.01g (Ohaus Scout SKX Precision Balance). One egg per clutch was randomly selected to be frozen at −80 °C, then later dissected for yolk hormone analysis (see details below) and the embryo staged according to number of somite pairs (Feiner et al., 2018) – providing a more precise measure of early embryonic stages than the staging scheme for lacertids (Dufaure & Hubert, 1961). The number of somite pairs of embryos at laying ranged from 20 (stage 25) to 33 (stage 27). Given the destructive nature of hormone analysis, we investigated the association between development time and egg yolk thyroid hormone concentration for different embryos within the same clutch (for incubation temperatures at 21 °C; 2e and 27 °C; Results Section 2f).

A recently developed high sensitivity LC-MS/MS method (Ruuskanen et al., 2018) was used to quantify the concentration of candidate thyroid hormones (TH) thyroxine (T4) and triiodothyronine (T3) in egg yolk samples. In brief, TH was extracted from 50mg of yolk per egg, and an internal tracer (a known amount of ^13^C_12_-T4, Larodan, Sweden) was added to control for variation in extraction efficiency between samples. Sample analysis was performed on a nanoflow HPLC system (Easy-nLC1000, Thermo Fisher Scientific) coupled to a triple quadrupole mass spectrometer (TSQ Vantage, Thermo Scientific, San Jose, CA) equipped with a nano-electrospray ionisation source. Two blanks (containing only plain reagent) were analysed at each extraction batch to control for contamination. Peak area ratios of samples relative to their internal standard (^13^C_6_-T3 and ^13^C_6_-T4, Sigma Aldrich) were calculated using Thermo Xcalibur software and Skyline, respectively (MacLean et al., 2010). For full details on sample and standard preparation and TH extraction see Ruuskanen et al., (2018).

### Embryo energy expenditure measures

Eggs were checked daily for signs of hatching, and development time (*D;* Results Section 3a) recorded as the number of days from oviposition until hatching. We measured the rate of carbon dioxide, CO_2_, production 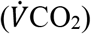 as a proxy for metabolic rate (*MR*) following standard closed-system, stop-flow respirometry as per (Cordero et al., 2017). Individual eggs were placed in a 60 ml glass chamber containing 50 ml moist vermiculite. Chambers were then flushed with incurrent air at 200 (±20) ml min^−1^ flow rate (standard temperature and pressure, STP). Sealed vials were then placed into incubators at the measurement temperature. After 180 min, the baseline CO_2_ was measured, and the chambers re-connected to the airflow (200±20 ml min^−1^, STP) and flushed. To measure CO_2_, the excurrent airflow (leaving the chamber) was scrubbed of water using drierite desiccant (Hammond Drierite) before entering a FC-10 CO_2_ analyser (Sable Systems International, Las Vegas, NV, USA). Using the metabR package (Noble, 2021), %CO_2_ was converted to 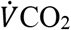 by integrating the area under the curve from the baseline over the three hour period to obtain 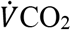 in ml h^−1^ (Lighton, 2018).

*MR* for embryos incubated under 21 °C or 27 °C was measured at stage 39 – approximately 75% of the total incubation time. The number of days it took to reach this stage for each region was first estimated from existing data (While et al., 2015) and confirmed by dissecting embryos (from first clutches) at different incubation times, accounting for both temperature treatment and region. We measured *MR* under one of two conditions: 1) *MR* under the same, single measurement temperature as the incubation treatment: cool (21 °C) or warm (27 °C) (Results Section 3b), or 2) *MR_T_*: across three temperatures (21 °C, 27 °C, and 33 °C; Results Section 3c). This design allowed us to measure embryos in two batches to determine 1) whether lower *D* necessitates higher *MR* by measuring these traits at a constant temperature for the same individuals, and 2) how region of origin and developmental temperature may influence metabolism thermal reaction norms (i.e., *MR_T_*) (see Experimental design; Figure 1B).

### Embryo energy allocation measures

After *MR_T_* was measured, embryos were euthanised and dissected to confirm developmental stage at 75% development (i.e., stage 39; Dufaure & Hubert (1961)), and sampled for yolk and tissue mass. The ratio of yolk mass relative to embryo mass gives an indication of the amount of energy reserves used for development in proportion to the amount of structure produced up to this stage of development (Pettersen et al., 2018). Yolk:tissue mass (Table 2, 3a) was quantified by separating, oven drying (60 °C for 3 days), and individually weighing embryo and yolk to the nearest 0.01mg (Sartorius Research 200D microbalance).

All hatchling trait measurements were carried out by AKP and TU. Upon hatching, individuals of known *MR* and *D* were measured in terms of hatchling mass (to the nearest 0.01g; 4b) and hatchling length (snout-to-vent length; SVL, to the nearest 0.5mm; Results Section 4c). Since embryo survival to hatching was high across both incubation temperatures (21 °C: 97%, 27 °C: 98%), this measure was not analysed.

### Statistical analyses

#### i. Overview

All analyses were conducted in R version 4.0.3 (R Core Team, 2020). For hypothesis tests, linear models were fitted using the “lm” function for the response variable of clutch number (Table 1; 1a) and yolk:embryo mass (Table 1; 4a). Hypothesis tests for all other responses (Table 1; 1-4) were analysed using linear mixed effects models (“lmer” function within the *lme4* package; (D. Bates et al., 2015). Post-hoc tests using the *emmeans* (Lenth, 2022) and *lmerTest* (Kuznetsova et al., 2017) packages were run to compare pairwise differences in factors within the final linear and linear mixed effects models, respectively.

**Table 1.**
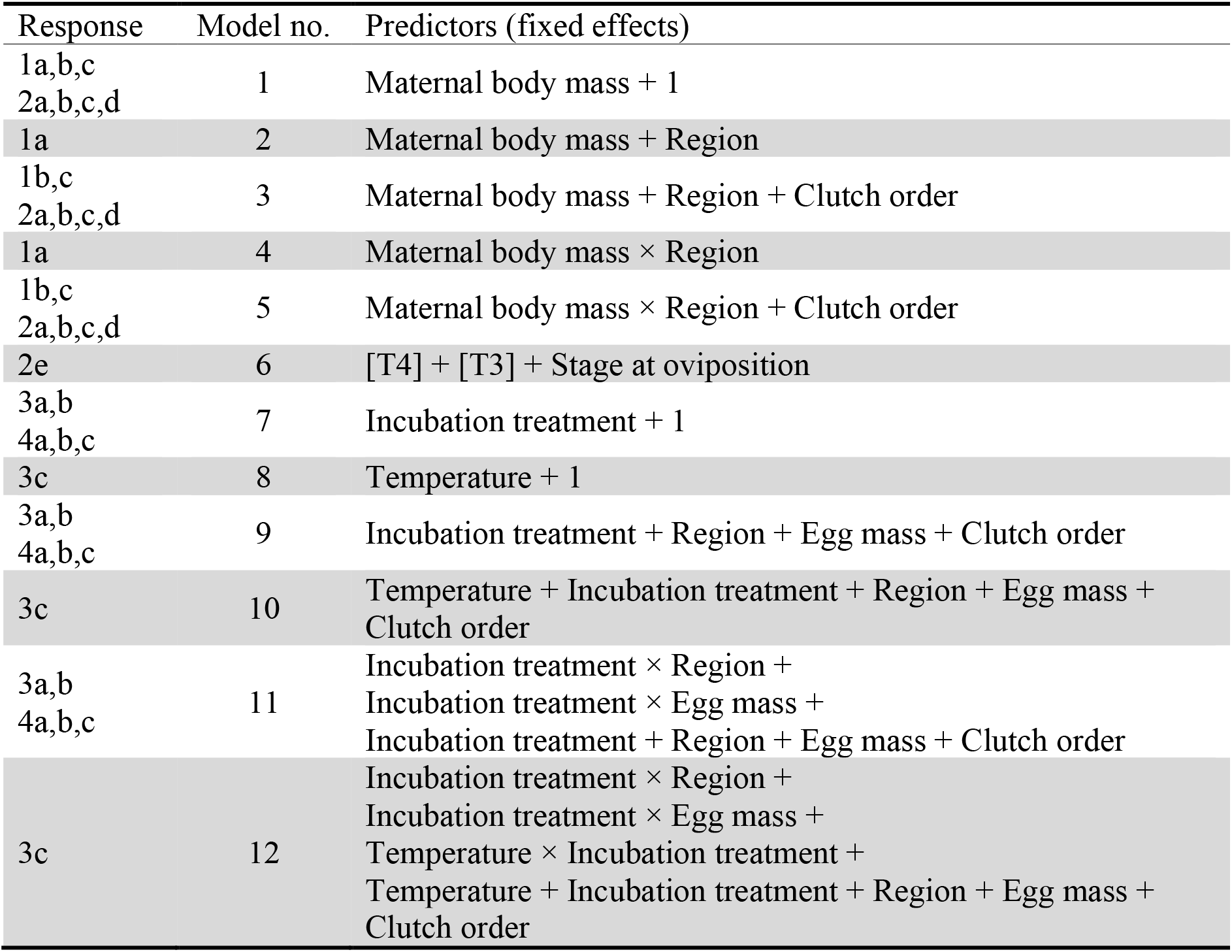
Summary of candidate models used in model selection for 1. Reproductive output (a. Clutch number, b. Clutch size, c. Clutch mass), 2. Maternal investment (a. Egg mass, b. Embryo retention, c. T4 thyroid hormone concentration, d. T3 thyroid hormone concentration, e. Development time), 3. Energy use (a. Development time, b. Metabolic rate (*MR*), c. Metabolic rate (*MR_T_*), 4. Energy allocation (a. Tissue:Total embryo mass, b. Hatchling mass, c. Hatchling length), and 5. Covariation between Development time and Metabolic rate with T4 and T3 Thyroid hormone concentration. Maternal ID was included as a random effect for all models except for 1a (Clutch number). For Metabolic rate thermal sensitivity (*MR_T_*; 3d), Embryo ID was included as a random effect in a repeated measures analysis.

| Response | Model no. | Predictors (fixed effects) |
| --- | --- | --- |
| 1a,b,c<br>2a,b,c,d | 1 | Maternal body mass + 1 |
| 1a | 2 | Maternal body mass + Region |
| 1b,c<br>2a,b,c,d | 3 | Maternal body mass + Region + Clutch order |
| 1a | 4 | Maternal body mass × Region |
| 1b,c<br>2a,b,c,d | 5 | Maternal body mass × Region + Clutch order |
| 2e | 6 | [T4] + [T3] + Stage at oviposition |
| 3a,b<br>4a,b,c | 7 | Incubation treatment + 1 |
| 3c | 8 | Temperature + 1 |
| 3a,b<br>4a,b,c | 9 | Incubation treatment + Region + Egg mass + Clutch order |
| 3c | 10 | Temperature + Incubation treatment + Region + Egg mass + Clutch order |
| 3a,b<br>4a,b,c | 11 | Incubation treatment × Region +<br>Incubation treatment × Egg mass +<br>Incubation treatment + Region + Egg mass + Clutch order |
| 3c | 12 | Incubation treatment × Region +<br>Incubation treatment × Egg mass +<br>Temperature × Incubation treatment +<br>Temperature + Incubation treatment + Region + Egg mass + Clutch order |

#### ii. Model selection

Candidate models (summarised in Table 1) were used to test the effect of fixed factors: ‘Region’ (High, Mid, Low), ‘Maternal body mass’, and ‘Clutch number’ (and ‘Region × Maternal body mass’ interactions) as well as the random effect of ‘Mother ID’ on (1) Reproductive output and (2) Maternal investment. Maternal body mass was included as a covariate to account for effects of body size on reproductive traits across altitudes, and stage at oviposition was included as a covariate for models testing the relationship between development time and yolk thyroid hormone concentration (Results Section 2c). For (3) Energy use and (4) Energy allocation analyses, ‘Incubation treatment’, ‘Region’, ‘Egg Mass’, ‘Clutch order’ and interactions between ‘Incubation treatment × Region’ and ‘Incubation treatment × Egg Mass’ were included as fixed factors. Where response data on more than one egg per clutch was collected, we used the average value per clutch to account for non-independence (Harrison et al., 2018). For linear mixed effects models testing the response of *MR* across ‘Temperature’ (21 °C, 27 °C, 33 °C) and its interaction with Incubation treatment (21 °C or 27 °C), ‘Embryo ID’ was treated as a random effect. We used Akaike Information Criterion (AIC) to rank candidate models and averaged those with Δ conditional Akaike information criterion (AICc) <4 using the R package *MuMin* v.1.43.17 (Barton, 2009).

#### iii. Predictor and response variable transformations

Box-cox transformations were applied to thyroid yolk hormone T4 and T3 concentrations prior to analysis (Results Section 2c – 2e) – a method often used for endocrine data to meet the assumptions of homoscedasticity and normality of model residuals (Miller & Plessow, 2013). Natural log transformations of egg mass and metabolic rate (for *MR*; 3b, and *MR_T_*; 3c models) were used for all hypothesis tests, to reduce observed variation with the mean and thereby satisfy the assumption of heteroscedasticity, and to allow for meaningful interpretation of scaling patterns (Glazier, 2021; Niklas & Hammond, 2014).

#### iv. Calculation of MR_T_

The relationship between metabolic rate and temperature was nonlinear for *MR_T_* (thermal reaction norm under three consecutive temperatures (i.e., 21 °C/27 °C/33 °C), we used the function “nls” in Base-R to obtain weighted least-squares parameter estimates of treatment and region-level thermal sensitivities (i.e., the slope) of *MR* (D. M. Bates & Chambers, 1992). We fitted separate exponential functions for each region and incubation treatment, whereby *MR* = *f* × exp^*a*×Temperature^.

## Results

### 1. Reproductive output

Mothers from Low-(mean_body mass_: 6.34 ± SD: 0.66 g) and Mid-(mean_body mass_: 6.07 ± SD: 1.21 g) altitude regions had larger body sizes than High-altitude region mothers (mean_body mass_: 4.64 ± SD: 0.62 g). Heavier females laid larger, heavier clutches, resulting in no significant effect of region on clutch characteristics after accounting for mother body mass (Table S4). Low-region females did however lay more clutches than High-region females (Table 2).

**Table 2.**
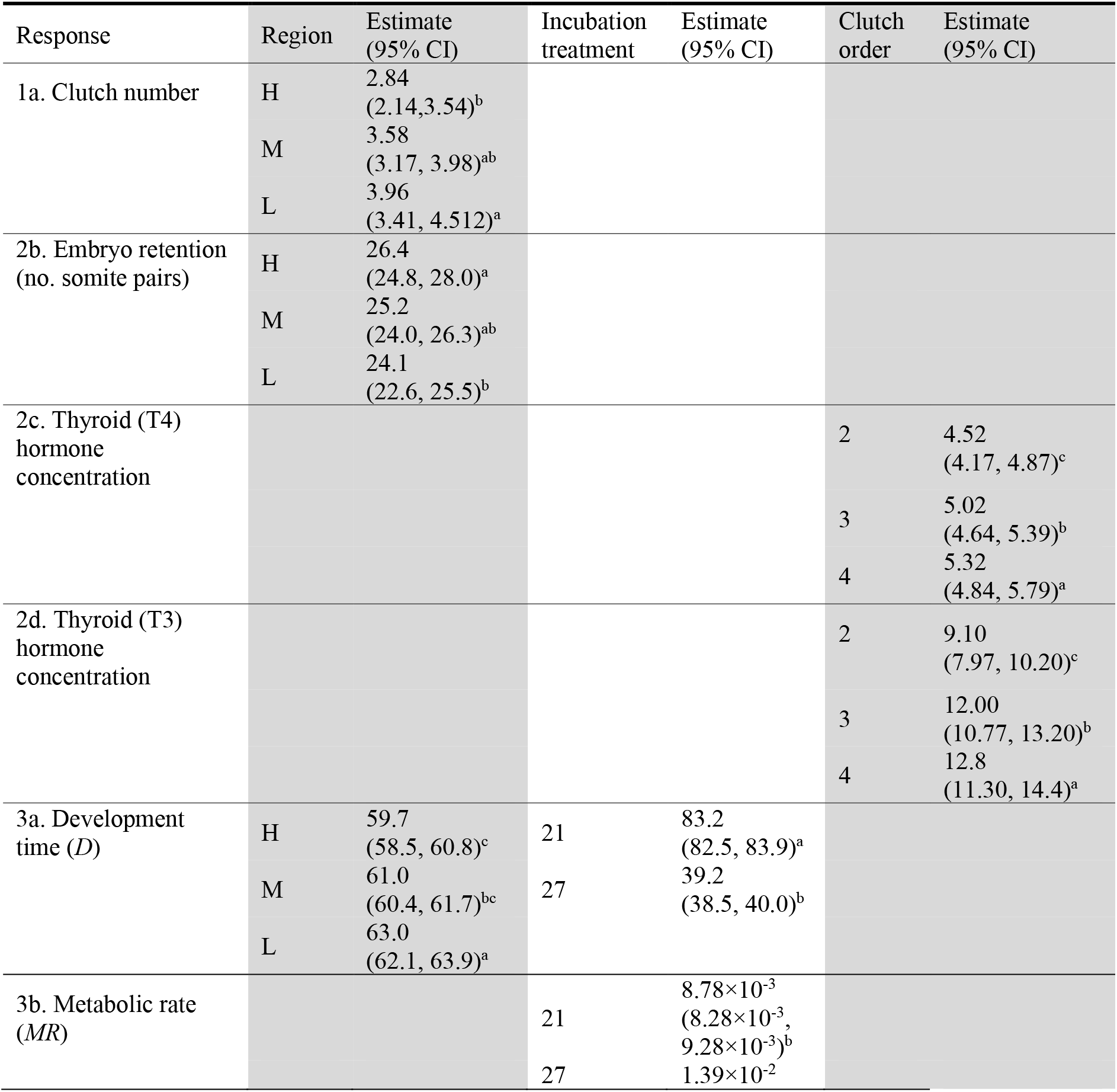
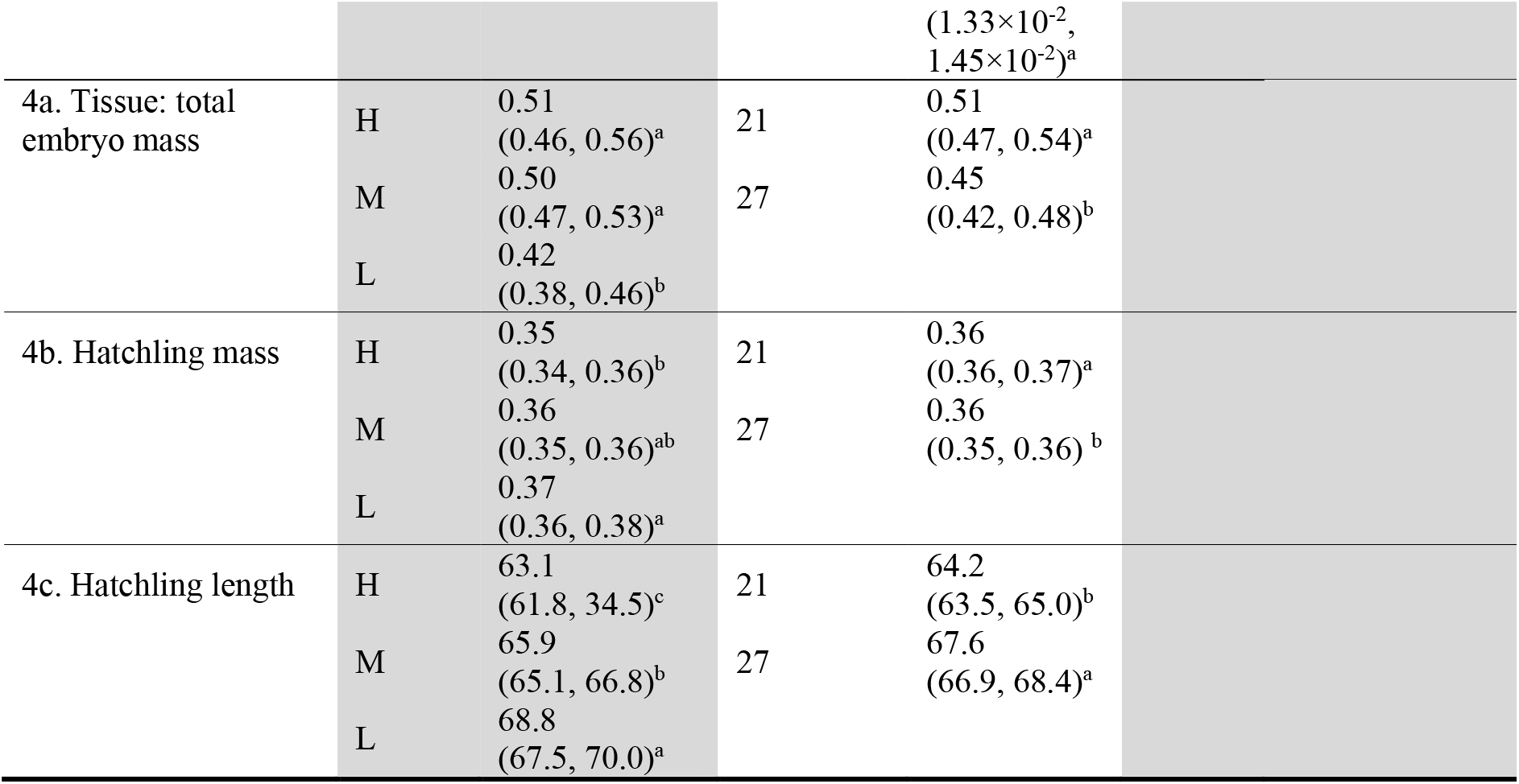
Post-hoc comparison of mean estimates for Region (H = High, M = Mid, L = Low), Incubation treatment (21 °C or 27 °C), and Clutch order (1^st^ – 4^th^) factors based on significant differences from best-fitting models (see Tables S3 and S4). Only responses containing significant differences in factors presented: 1. Reproductive output (a. Clutch number, b. Clutch size, c. Clutch mass), 2. Maternal investment (b. Embryo retention, c. Thyroid hormone (T4) concentration, d. Thyroid hormone (T3) concentration), 3. Embryo energy use (a. Development time, e. Yolk:embryo mass), and 4. Embryo energy allocation (a. Hatchling mass, b. Hatchling length). Letters indicate significant differences among groups (a>b>c>d).

| Response | Region | Estimate<br>(95% CI) | Incubation<br>treatment | Estimate<br>(95% CI) | Clutch<br>order | Estimate<br>(95% CI) |
| --- | --- | --- | --- | --- | --- | --- |
| 1a. Clutch number | H | 2.84<br>(2.14, 3.54)<sup>b</sup> |  |  |  |  |
|  | M | 3.58<br>(3.17, 3.98)<sup>ab</sup> |  |  |  |  |
|  | L | 3.96<br>(3.41, 4.512)<sup>a</sup> |  |  |  |  |
| 2b. Embryo retention<br>(no. somite pairs) | H | 26.4<br>(24.8, 28.0)<sup>a</sup> |  |  |  |  |
|  | M | 25.2<br>(24.0, 26.3)<sup>ab</sup> |  |  |  |  |
|  | L | 24.1<br>(22.6, 25.5)<sup>b</sup> |  |  |  |  |
| 2c. Thyroid (T4)<br>hormone<br>concentration |  |  |  |  | 2 | 4.52<br>(4.17, 4.87)<sup>c</sup> |
|  |  |  |  |  | 3 | 5.02<br>(4.64, 5.39)<sup>b</sup> |
|  |  |  |  |  | 4 | 5.32<br>(4.84, 5.79)<sup>a</sup> |
| 2d. Thyroid (T3)<br>hormone<br>concentration |  |  |  |  | 2 | 9.10<br>(7.97, 10.20)<sup>c</sup> |
|  |  |  |  |  | 3 | 12.00<br>(10.77, 13.20)<sup>b</sup> |
|  |  |  |  |  | 4 | 12.8<br>(11.30, 14.4)<sup>a</sup> |
| 3a. Development<br>time (*D*) | H | 59.7<br>(58.5, 60.8)<sup>c</sup> | 21 | 83.2<br>(82.5, 83.9)<sup>a</sup> |  |  |
|  | M | 61.0<br>(60.4, 61.7)<sup>bc</sup> | 27 | 39.2<br>(38.5, 40.0)<sup>b</sup> |  |  |
|  | L | 63.0<br>(62.1, 63.9)<sup>a</sup> |  |  |  |  |
| 3b. Metabolic rate<br>(*MR*) |  |  | 21 | 8.78×10<sup>-3</sup><br>(8.28×10<sup>-3</sup>,<br>9.28×10<sup>-3</sup>)<sup>b</sup> |  |  |
|  |  |  | 27 | 1.39×10<sup>-2</sup> |  |  |

|  |  |  |  |  |  |
| --- | --- | --- | --- | --- | --- |
| | | | | ($1.33 \times 10^{-2}$,<br>$1.45 \times 10^{-2}$)<sup>a</sup> | |
| 4a. Tissue: total embryo mass | H | 0.51<br>(0.46, 0.56)<sup>a</sup> | 21 | 0.51<br>(0.47, 0.54)<sup>a</sup> |  |
|  | M | 0.50<br>(0.47, 0.53)<sup>a</sup> | 27 | 0.45<br>(0.42, 0.48)<sup>b</sup> |  |
|  | L | 0.42<br>(0.38, 0.46)<sup>b</sup> |  |  |  |
| 4b. Hatchling mass | H | 0.35<br>(0.34, 0.36)<sup>b</sup> | 21 | 0.36<br>(0.36, 0.37)<sup>a</sup> |  |
|  | M | 0.36<br>(0.35, 0.36)<sup>ab</sup> | 27 | 0.36<br>(0.35, 0.36)<sup>b</sup> |  |
|  | L | 0.37<br>(0.36, 0.38)<sup>a</sup> |  |  |  |
| 4c. Hatchling length | H | 63.1<br>(61.8, 34.5)<sup>c</sup> | 21 | 64.2<br>(63.5, 65.0)<sup>b</sup> |  |
|  | M | 65.9<br>(65.1, 66.8)<sup>b</sup> | 27 | 67.6<br>(66.9, 68.4)<sup>a</sup> |  |
|  | L | 68.8<br>(67.5, 70.0)<sup>a</sup> |  |  |  |

### 2. Maternal investment

On average, mothers from High-altitude regions laid lighter eggs (mean_egg mass_: 0.28 ± SD: 0.03 g), compared with mothers from Mid-(0.29 ± 0.04 g) and Low-(0.30 ± 0.03 g) altitude regions (Figure 2A). However, after accounting for maternal body size, the effect of region was not statistically significant (Table S4). High-altitude region females laid embryos at slightly, but significantly later stages of development than mothers from the Low-altitude region, with an average difference of 2.3 somite pairs between High- and Low-altitude region embryos (Table 2, Figure 2B). There was a trend for higher [TH4] in eggs from Low-altitude mothers compared with other regions, however we found no statistically significant differences in either [TH4] or [TH3] among regions (Table 2; Figure 2C). Both forms of thyroid hormone increased in concentration from clutches 2 to 4 (Figure 2C). We found a significant interaction between Region and maternal body mass, where [TH4] decreased with female body size only in the eggs of Low-altitude Region mothers (Section 2c in Table S4). We found no association between development time and yolk thyroid hormones ([TH4] and [TH3]), and under either incubation temperature (Section 2e in Table S4).

**Figure 2.**
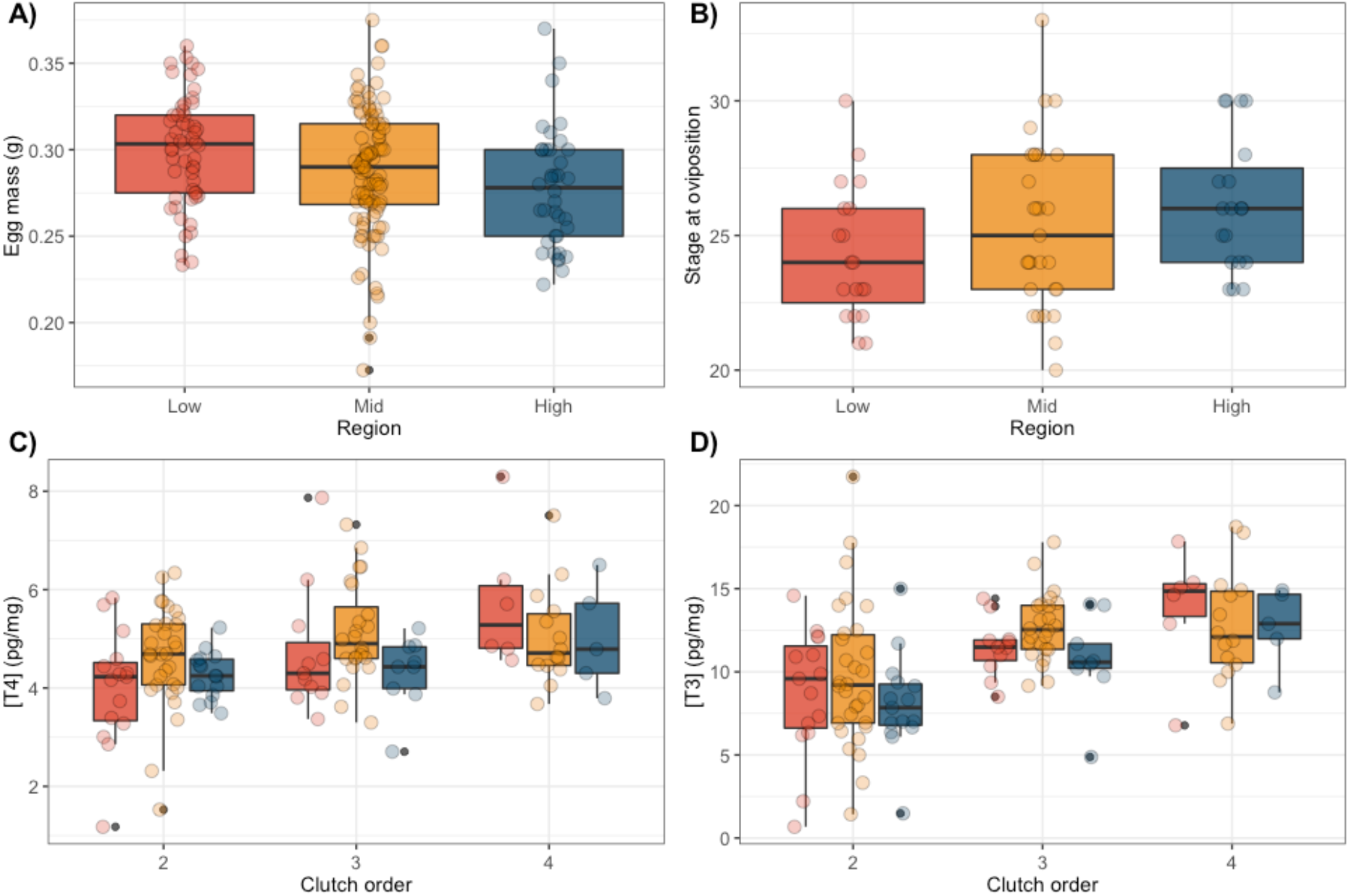
Maternal investment across regions and clutch order. Boxplots showing the distribution of data for A) egg mass (g), B) developmental stage at oviposition (number of somites) as a measure of the extent of embryo retention, 2C) T4 yolk thyroid hormone concentration (pg/mg; box-cox transformed), 2D) T3 yolk thyroid hormone concentration (pg/mg; box-cox transformed), for embryos from low (red), mid (orange), and high (blue) altitude regions and across 2^nd^ to 4^th^ clutch order (2C, 2D). Data points show raw data, coloured by embryo altitude region of origin (red = low, orange = mid, blue = high).

### 3. Embryo energy expenditure

#### a. Development time

Embryos incubated at 27 °C took less than half the number of days (mean 39.5 ± SD: 1.3) to hatch compared with those incubated at 21 °C (mean: 83.4 ± 4.9 days) (Table 2). Development time (*D*) was shorter for high- and mid-altitude embryos across both treatments, where high-altitude embryos hatched on average 3.7 days and 1.5 days earlier than low-altitude embryos at 21 °C and 27 °C, respectively (Table 2, Figure 3A). Development time decreased with egg mass (Table S4, 3a). There was a borderline non-significant effect of clutch order on development time, where subsequent clutches developed faster than the 2^nd^ clutch (Table S4: p = 0.06).

**Figure 3.**
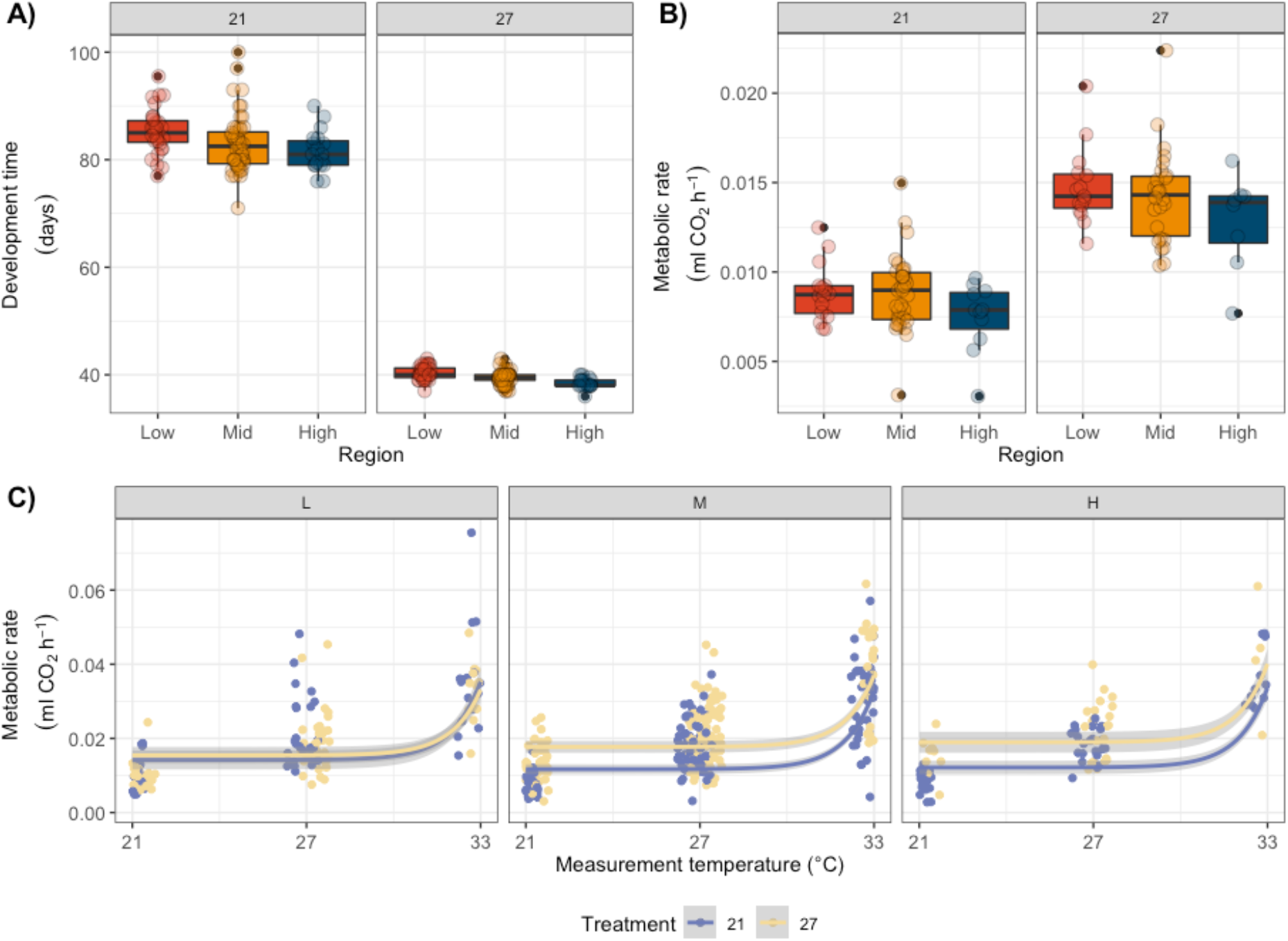
Embryo energy use across regions, incubation treatment, and measurement temperature. Boxplots showing the distribution of data for A) development time (days from oviposition to hatching), and B) metabolic rate (measured as rate of CO_2_ production; ml CO_2_ h^−1^), across 21 °C and 27 °C incubation treatments at stage 39 (75% through embryonic development). C) metabolic rate thermal reaction norms for embryos from Low (L), Mid (M), and High (H)-altitude regions when incubated at 21 °C (yellow) or 27 °C (purple) incubation treatments and measured at three temperatures (21 °C, 27 °C, 33 °C) at stage 39 (75% through embryonic development). Data points show raw data.

#### b. Metabolic rates

Embryos incubated and measured at 27 °C showed significantly higher metabolic rates than embryos incubated and measured at 21 °C (Table 2, Figure 3B). Incubation temperature also produced significant effects on the thermal sensitivity of metabolism (i.e., *MR_T_*). There was a significant interaction between incubation temperature and measurement temperature (Section 3c in Table S4: p<0.001), hence separate treatment slopes were fitted. The slope of *MR_T_* across three measurement temperatures (21 °C, 27 °C, 33 °C) was consistently steeper in embryos incubated at 21 °C than 27 °C. Furthermore, within the 21 °C treatment, embryos from High and Mid-altitude regions showed higher thermal sensitivity in *MR*, with steeper *MR_T_* slopes, compared with Low-altitude source regions (Table S5; Figure 3C).

### 4. Embryo energy allocation

Embryos from larger eggs hatched both heavier and longer (Table S4). At 75% of the total development (stage 39), embryos incubated at 21 °C produced a higher tissue:total embryo mass ratio than those incubated at 27 °C (Figure 4A, Table S4, 4a). Embryos from high and mid-regions also converted proportionally more of their yolk reserves into tissue, compared with low region embryos (Table 2). Despite this, Low-altitude region embryos hatched longer (and therefore heavier) than high-altitude region embryos (Figures 4B and 4C, Table 2). In line with embryos dissected at stage 39, incubation at 21 °C produced heavier and shorter hatchlings with a higher proportion of tissue mass relative to total mass, compared with incubation at 27 °C (Table 2).

**Figure 4.**
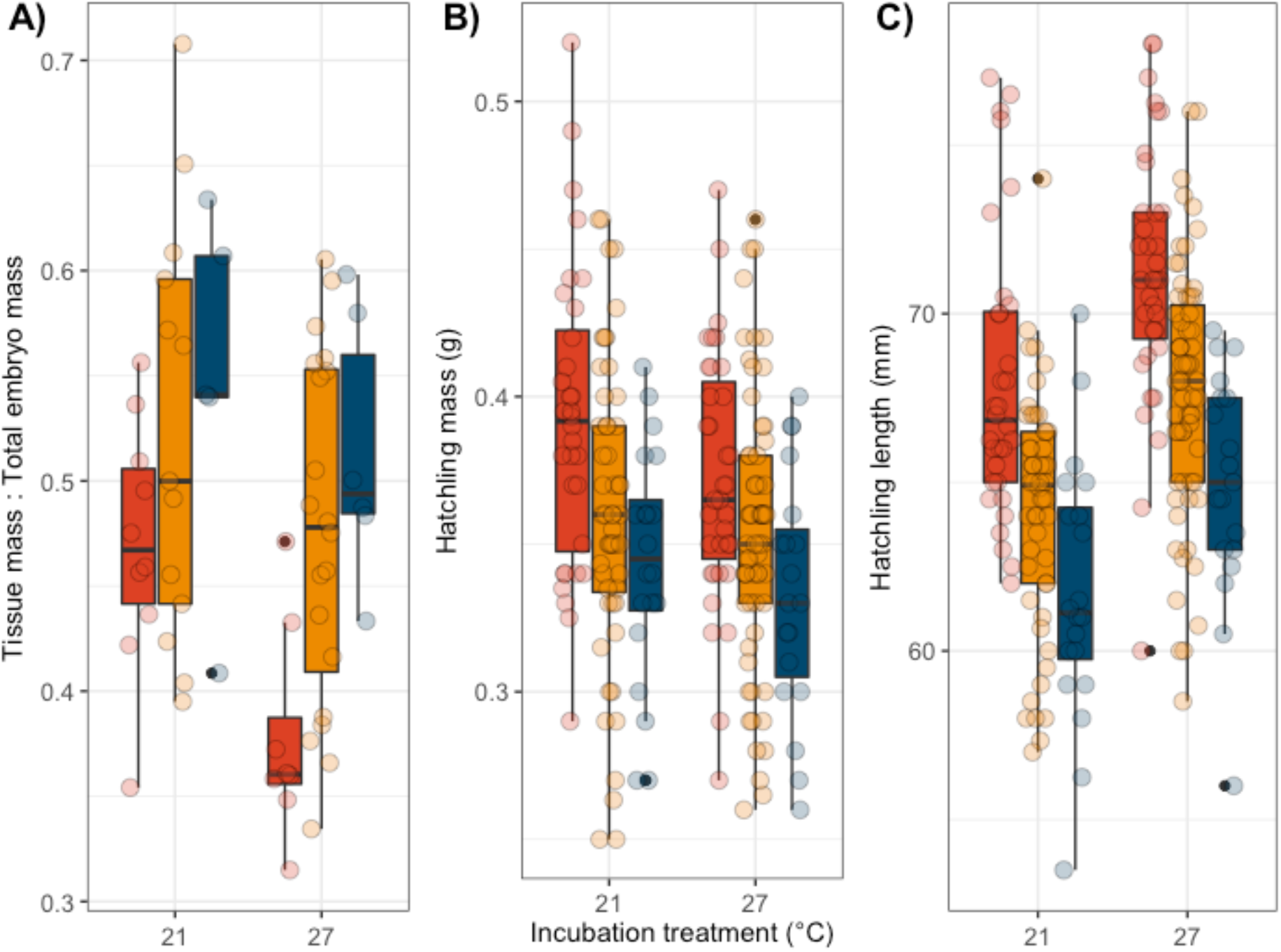
Embryo energy allocation across regions and incubation treatments. Boxplots showing the distribution of data for A) tissue mass:total embryo mass ratio at 75% development (stage 39), B) hatchling mass, and C) hatchling length from Low (red), Mid (orange), and High (blue)-altitude regions when incubated at 21 °C or 27 °C incubation treatments. Data points show average values per clutch.

## Discussion

Embryonic development can be a crucial bottleneck for individual survival, and therefore population range expansion, in cool climates. Partly this is due to the greater energy cost (‘cost of development’) associated with low incubation temperatures since cool temperatures tend to increase the overall energy required to complete development (Pettersen et al., 2019). This cost may be offset by adaptive changes in investment from mothers that reduce the time spent in the egg, or via shifts in embryo energy expenditure or allocation from yolk towards tissue (Cook et al., 2018; Mueller et al., 2015; Sinervo & Huey, 1990). In this experiment, we established the relative importance of these three potential mechanisms across three regions of wall lizards locally adapted to different altitudinal gradients.

### Maternal investment does not compensate for increased cost of development

As anticipated from comparative data (Pettersen et al., 2019), the energy expended to complete embryonic development was greater at cool relative to warm incubation temperatures. Mothers can compensate for this cost in several ways. Previous studies of reptiles have demonstrated extended embryo retention in egg-laying species under cooler climates (Rodríguez-Díaz & Braña, 2012; Shine, 1995), including for wall lizard populations introduced to England from southern France and Italy (While et al., 2015). In the present study, population differences in embryo retention were statistically significant but biologically rather minor. This variation could arguably be explained by embryos from high-altitude populations developing faster at the preferred body temperature of gravid females (which is the same across the three regions; Ruiz Miñano (2021)), without the need for any change in female reproductive physiology *per se*.

Rather than reducing the cost of development by facilitating earlier hatching dates, mothers may instead help to compensate for costly development at cool temperatures by producing larger eggs with greater energy reserves (Pettersen et al., 2019). We found no evidence for cool-climate, high-altitude region females investing more per offspring by laying heavier eggs, although they did lay fewer clutches across the season. The latter is consistent with previous work on this species (MacGregor et al. (2017)) and is likely because the shorter season at high altitude has favoured stricter termination of the reproductive cycle (Le Henanff et al., 2013; While et al., 2015).

A third mechanism of maternal investment is by modifying the content of the yolk to accelerate development of their offspring (T. G. Groothuis et al., 2020). Yolk thyroid hormones T4 and T3 are major determinants of metabolic rate and energy metabolism in vertebrates, and have been highlighted as potentially important mediators underlying maternal effects across vertebrate taxa (Ruuskanen & Hsu, 2018; Sarraude et al., 2020) and thyroid hormone concentration has been found to be correlated with development time in birds (Hsu et al., 2022). However, since there were no significant differences in either T4 or T3 concentration along the altitudinal gradient studied, it is unlikely that variation in yolk thyroid hormones can explain why cool-adapted lizard embryos developed faster than warm-adapted embryos. Despite a lack of evidence for any regional differences, we did find substantial among-clutch variation in T3 and T4 concentration, in line with previous work on birds (Hsu et al., 2019). With each subsequent clutch laid in captivity, mothers increased yolk T4 and T3 concentration. Later stage clutches (3^rd^ and 4^th^) were also found to develop faster than previous clutches, although we found no direct association between development time and T4 or T3 concentration. Thus, maternal thyroid hormone does not seem to influence rate of development or growth in wall lizard embryos.

### Region-level differences in embryo energy expenditure during development

Given our observation of overall weak maternal effects (embryo retention, egg mass, and yolk thyroid hormone), it may be that the evolution of faster embryonic developmental rates, both before and after oviposition, are more important for reducing the cost of development and early-life survival at cool temperatures in reptiles (Du et al., 2019). Indeed, we found that wall lizard embryos from a relatively cool region developed faster compared with embryos from warmer source populations, without any apparent increase in metabolic rate. These results provide evidence that the evolution of developmental and metabolic rates can be decoupled (as shown previously for growth and metabolic rates; Williams et al., (2016)).

We found support for regional differences in the thermal sensitivity of developmental, and to a lesser extent metabolic, rates demonstrating that the thermal sensitivity of these traits does evolve within species. We also found evidence for regional differences in the temperature sensitivity across incubation treatments (also known as thermal developmental plasticity); embryos incubated at 21 °C showed greater sensitivity in their metabolic rates to increasing measurement temperatures (21 °C to 33 °C), compared with those reared at 27 °C. Developmental temperatures can produce variation in the plasticity of physiological traits to affect post-hatching growth and survival, and reproductive success (Andrews et al., 2000; Kang et al., 2021; Kar et al., 2022; Nord & Nilsson, 2011). Greater thermal sensitivity in metabolic rates in response to cool nest temperatures will likely serve an advantage, enabling individuals to make better use of warmer conditions when they do arise, and thereby hatch earlier, grow faster, and reach a size refuge (i.e., minimum body size) from predation sooner. However, whether the difference between populations reflects selection for greater plasticity in cooler developmental temperatures, and if it enables the expansion of egg laying ectotherms into cool climates needs further examination (Carbonell et al., 2021).

### Allocation of egg yolk reserve relative to embryo structure

Lizards often hatch with residual yolk which is assimilated in the juvenile stage, and therefore thought to serve a functional role in post-hatching survival (Radder et al., 2007). A third response to the increased cost of development at low incubation temperatures is therefore to maintain conversion to tissue during embryonic development at the expense of residual yolk. Alternatively, if residual yolk is more important, offspring may hatch relatively smaller for a given egg size when incubated at cool temperatures. At 75% completion of development, we found wall lizard embryos incubated at 21 °C possessed a higher proportion of tissue relative to total mass, compared with those incubated at 27 °C – an observation that is in line with previous work showing a positive effect of cooler incubation conditions on hatchling size in reptiles (Janzen et al., 1990; Packard et al., 1988). The shift towards greater tissue production at the expense of residual energy reserves at cool temperatures may explain the observation that offspring often hatch larger, despite a higher cost of development, in cool temperatures (Ji & Du, 2001; Pettersen et al., 2019). When comparing regions, our hypothesis that embryos from high altitudes utilise more yolk reserves to produce more tissue was supported. It may be that higher metabolic rates and therefore higher maintenance and growth costs in warmer temperatures favour greater residual yolk at hatching for low-altitude regions (Barneche et al., 2019; Liess et al., 2013), yet further investigation is needed to determine whether this is indeed an adaptive response and how it has evolved.

In summary, this study finds that cool temperatures, characteristic of those experienced by wall lizards at high altitudes, exacerbate the energy required to successfully complete hatching. To compensate for this energy cost, high-altitude embryos increase developmental rate without a concomitant increase in metabolic rate, and therefore expend less energy completing development at one and the same incubation temperature than embryos from lower altitudes. The proximal mechanisms underlying this response are unclear, however the enhanced allocation of energy reserves towards tissue observed in high altitude embryos provides an indication of the potential regulatory drivers that enable local adaptation of egg-laying ectotherms to cool climates.

## Acknowledgements

The authors wish to thank Fonti Kar for assistance with the *metabR* package, and Eric Gangloff and Gerardo Antonio Cordero for advice on stop-flow analysis. This work was funded by a Wenner-Gren Foundation Postdoctoral Fellowship to AKP and TU (UPD2019-0208) and a research grant from the Swedish Research Council to TU (2017-03846).

## Author Contributions

AKP and TU conceived the study design. AKP, JFN, MRM, LJF, GMW and TU contributed to data collection, and AKP, AN, SR and TU analysed and interpreted the data. AKP wrote the initial draft, with revisions from AN and TU. All authors contributed critically to the initial version and gave final approval for publication.

## Data Availability Statement

All data and code have been made available for peer review on the Open Science Framework: DOI 10.17605/OSF.IO/SRKBJ.

## References

Andrews, R. M., Mathies, T., & Warner, D. A. (2000). Effect of Incubation Temperature on Morphology, Growth, and Survival of Juvenile Sceloporus undulatus. Herpetological Monographs, 14, 420–431. JSTOR. https://doi.org/10.2307/1467055

Atkinson, D. (1996). Ectotherm life-history responses to developmental temperature. In A. F. Bennett & I. A. Johnston (Eds.), Animals and Temperature: Phenotypic and Evolutionary Adaptation (pp. 183–204). Cambridge University Press. https://doi.org/10.1017/CBO9780511721854.009

Atkinson, D., Morley, S. A., Weetman, D., & Hughes, R. N. (2001). Offspring size responses to maternal temperature in ectotherms. Environment and Animal Development: Genes, Life Histories and Plasticity, 269, 85.

Badyaev, A. V. (2009). Evolutionary significance of phenotypic accommodation in novel environments: An empirical test of the Baldwin effect. Philosophical Transactions of the Royal Society B-Biological Sciences, 364(1520), Article 1520. https://doi.org/10.1098/rstb.2008.0285

Barneche, D. R., Jahn, M., & Seebacher, F. (2019). Warming increases the cost of growth in a model vertebrate. Functional Ecology, 33(7), Article 7. https://doi.org/10.1111/1365-2435.13348

Barton, K. (2009). MuMIn: Multi-model inference. *http://R-Forge.r-Project.Org/Projects/Mumin/*. https://cir.nii.ac.jp/crid/1572824499154168192

Bates, D. M., & Chambers, J. M. (1992). Chapter 10: Nonlinear Models. In Statistical Models in S.

Bates, D., Mächler, M., Bolker, B., & Walker, S. (2015). Fitting Linear Mixed-Effects Models Using lme4. Journal of Statistical Software, 67(1), Article 1. https://doi.org/10.18637/jss.v067.i01

Bownds, C., Wilson, R., & Marshall, D. J. (2010). Why do colder mothers produce larger eggs? An optimality approach. The Journal of Experimental Biology, 213(Pt 22), 3796–3801. https://doi.org/10.1242/jeb.043356

Braña, F., & Ji, X. (2000). Influence of incubation temperature on morphology, locomotor performance, and early growth of hatchling wall lizards (Podarcis muralis). The Journal of Experimental Zoology, 286(4), 422–433. https://doi.org/10.1002/(sici)1097-010x(20000301)286:4<422::aid-jez10>3.0.co;2-d

Burgess, S. C., & Marshall, D. J. (2011). Temperature-induced maternal effects and environmental predictability. Journal of Experimental Biology, 214(14), Article 14. https://doi.org/10.1242/jeb.054718

Calosi, P., Bilton, D. T., Spicer, J. I., Votier, S. C., & Atfield, A. (2010). What determines a species’ geographical range? Thermal biology and latitudinal range size relationships in European diving beetles (Coleoptera: Dytiscidae). Journal of Animal Ecology, 79(1), 194–204. https://doi.org/10.1111/j.1365-2656.2009.01611.x

Carbonell, J. A., Wang, Y.-J., & Stoks, R. (2021). Evolution of cold tolerance and thermal plasticity in life history, behaviour and physiology during a poleward range expansion. Journal of Animal Ecology, 90(7), 1666–1677. https://doi.org/10.1111/1365-2656.13482

Conover, D. O., & Present, T. M. C. (1990). Countergradient Variation in Growth-Rate— Compensation for Length of the Growing-Season Among Atlantic Silversides from Different Latitudes. Oecologia, 83(3), Article 3. https://doi.org/10.1007/BF00317554

Cook, C. J., Burness, G., & Wilson, C. C. (2018). Metabolic rates of embryos and alevin from a cold-adapted salmonid differ with temperature, population and family of origin: Implications for coping with climate change. Conservation Physiology, 6(1). https://doi.org/10.1093/conphys/cox076

Cordero, G. A., Andersson, B. A., Souchet, J., Micheli, G., Noble, D. W. A., Gangloff, E. J., Uller, T., & Aubret, F. (2017). Physiological plasticity in lizard embryos exposed to high-altitude hypoxia. Journal of Experimental Zoology Part A: Ecological and Integrative Physiology, 327(7), Article 7. https://doi.org/10.1002/jez.2115

Crozier, L. (2003). Winter warming facilitates range expansion: Cold tolerance of the butterfly Atalopedes campestris. Oecologia, 135(4), 648–656. https://doi.org/10.1007/s00442-003-1219-2

Du, W.-G., Shine, R., Ma, L., & Sun, B.-J. (2019). Adaptive responses of the embryos of birds and reptiles to spatial and temporal variations in nest temperatures. Proceedings of the Royal Society B: Biological Sciences, 286(1915), Article 1915. https://doi.org/10.1098/rspb.2019.2078

Du, W.-G., Zhao, B., Chen, Y., & Shine, R. (2011). Behavioral thermoregulation by turtle embryos. Proceedings of the National Academy of Sciences of the United States of America, 108(23), 9513–9515. https://doi.org/10.1073/pnas.1102965108

Dufaure, J. P., & Hubert, J. (1961). Table de developpement du lézard vivipare: Lacerta (Zootoca) vivipara Jacquin. Arch Anat Microscop. Morphol Exp, 50, 309–328.

DuRant, S. E., Hopkins, W. A., & Hepp, G. R. (2011). Embryonic Developmental Patterns and Energy Expenditure Are Affected by Incubation Temperature in Wood Ducks (Aix sponsa). Physiological and Biochemical Zoology, 84(5), 451–457. https://doi.org/10.1086/661749

Engelstoft, C., Robinson, J., Fraser, D., & Hanke, G. (2020). Recent rapid expansion of common wall lizards (Podarcis muralis) in British Columbia, Canada. Northwestern Naturalist, 101(1), 50–55. https://doi.org/10.1898/1051-1733-101.1.50

Feiner, N., Rago, A., While, G. M., & Uller, T. (2018). Signatures of selection in embryonic transcriptomes of lizards adapting in parallel to cool climate. Evolution, 72(1), 67–81. https://doi.org/10.1111/evo.13397

Glazier, D. S. (2021). Biological scaling analyses are more than statistical line fitting. Journal of Experimental Biology, 224(11). https://doi.org/10.1242/jeb.241059

Groothuis, T. G. G., & Schwabl, H. (2008). Hormone-mediated maternal effects in birds: Mechanisms matter but what do we know of them? Philosophical Transactions of the Royal Society B-Biological Sciences, 363(1497), Article 1497. https://doi.org/10.1098/rstb.2007.0007

Groothuis, T. G., Kumar, N., & Hsu, B.-Y. (2020). Explaining discrepancies in the study of maternal effects: The role of context and embryo. Current Opinion in Behavioral Sciences, 36, 185–192. https://doi.org/10.1016/j.cobeha.2020.10.006

Hamdoun, A., & Epel, D. (2007). Embryo stability and vulnerability in an always changing world. Proceedings of the National Academy of Sciences, 104(6), 1745–1750. https://doi.org/10.1073/pnas.0610108104

Harrison, X. A., Donaldson, L., Correa-Cano, M. E., Evans, J., Fisher, D. N., Goodwin, C. E. D., Robinson, B. S., Hodgson, D. J., & Inger, R. (2018). A brief introduction to mixed effects modelling and multi-model inference in ecology. PeerJ, 6, e4794. https://doi.org/10.7717/peerj.4794

Hsu, B.-Y., Pakanen, V.-M., Boner, W., Doligez, B., Eeva, T., Groothuis, T. G. G., Korpimäki, E., Laaksonen, T., Lelono, A., Monaghan, P., Sarraude, T., Thomson, R. L., Tolvanen, J., Tschirren, B., Vásquez, R. A., & Ruuskanen, S. (2022). Maternally transferred thyroid hormones and life-history variation in birds. Journal of Animal Ecology, 91(7), 1489–1506. https://doi.org/10.1111/1365-2656.13708

Hsu, B.-Y., Verhagen, I., Gienapp, P., Darras, V. M., Visser, M. E., & Ruuskanen, S. (2019). Between- and Within-Individual Variation of Maternal Thyroid Hormone Deposition in Wild Great Tits (Parus major). The American Naturalist, 194(4), E96–E108. https://doi.org/10.1086/704738

Janzen, F. J., Packard, G. C., Packard, M. J., Boardman, T. J., & Zumbrunnen, J. R. (1990). Mobilization of lipid and protein by embryonic snapping turtles in wet and dry environments. Journal of Experimental Zoology, 255(2), 155–162. https://doi.org/10.1002/jez.1402550204

Ji, X., & Braña, F. (1999). The influence of thermal and hydric environments on embryonic use of energy and nutrients, and hatchling traits, in the wall lizards (Podarcis muralis). Comparative Biochemistry and Physiology Part A: Molecular & Integrative Physiology, 124(2), Article 2. https://doi.org/10.1016/S1095-6433(99)00111-7

Ji, X., & Du, W. G. (2001). The effects of thermal and hydric environments on hatching success, embryonic use of energy and hatchling traits in a colubrid snake, Elaphe carinata. Comparative Biochemistry and Physiology. Part A, Molecular & Integrative Physiology, 129(2–3), Article 2–3. https://doi.org/10.1016/s1095-6433(01)00271-9

Kang, C.-Q., Meng, Q.-Y., Dang, W., & Lu, H.-L. (2021). Divergent incubation temperature effects on thermal sensitivity of hatchling performance in two different latitudinal populations of an invasive turtle. Journal of Thermal Biology, 100, 103079. https://doi.org/10.1016/j.jtherbio.2021.103079

Kar, F., Nakagawa, S., & Noble, D. W. A. (2022). Impact of developmental temperatures on thermal plasticity and repeatability of metabolic rate. Evolutionary Ecology, 36(2), 199–216. https://doi.org/10.1007/s10682-022-10160-1

Kuznetsova, A., Brockhoff, P. B., & Christensen, R. H. B. (2017). **lmerTest** Package: Tests in Linear Mixed Effects Models. Journal of Statistical Software, 82(13). https://doi.org/10.18637/jss.v082.i13

Laugen, A. T., Laurila, A., Räsänen, K., & Merilä, J. (2003). Latitudinal countergradient variation in the common frog (Rana temporaria) development rates – evidence for local adaptation. Journal of Evolutionary Biology, 16(5), Article 5. https://doi.org/10.1046/j.1420-9101.2003.00560.x

Le Henanff, M., Meylan, S., & Lourdais, O. (2013). The sooner the better: Reproductive phenology drives ontogenetic trajectories in a temperate squamate (Podarcis muralis). Biological Journal of the Linnean Society, 108(2), Article 2. https://doi.org/10.1111/j.1095-8312.2012.02005.x

Lembrechts, J. J., van den Hoogen, J., Aalto, J., Ashcroft, M. B., De Frenne, P., Kemppinen, J., Kopecký, M., Luoto, M., Maclean, I. M. D., Crowther, T. W., Bailey, J. J., Haesen, S., Klinges, D. H., Niittynen, P., Scheffers, B. R., Van Meerbeek, K., Aartsma, P., Abdalaze, O., Abedi, M., … Lenoir, J. (2022). Global maps of soil temperature. Global Change Biology, 28(9), 3110–3144. https://doi.org/10.1111/gcb.16060

Lenth, R. V. (2022). Estimated Marginal Means, aka Least-Squares Means (R package version 1.7.4-1).

Liess, A., Rowe, O., Guo, J., Thomsson, G., & Lind, M. I. (2013). Hot tadpoles from cold environments need more nutrients – life history and stoichiometry reflects latitudinal adaptation. Journal of Animal Ecology, 82(6), 1316–1325. https://doi.org/10.1111/1365-2656.12107

Lighton, J. R. B. (2018). Measuring Metabolic Rates: A Manual for Scientists. Oxford University Press.

MacGregor, H. E. A., While, G. M., & Uller, T. (2017). Comparison of reproductive investment in native and non-native populations of common wall lizards reveals sex differences in adaptive potential. Oikos, 126(11), 1564–1574. https://doi.org/10.1111/oik.03984

MacLean, B., Tomazela, D. M., Shulman, N., Chambers, M., Finney, G. L., Frewen, B., Kern, R., Tabb, D. L., Liebler, D. C., & MacCoss, M. J. (2010). Skyline: An open source document editor for creating and analyzing targeted proteomics experiments. Bioinformatics, 26(7), 966–968. https://doi.org/10.1093/bioinformatics/btq054

Marshall, D. J., Pettersen, A. K., Bode, M., & White, C. R. (2020). Developmental cost theory predicts thermal environment and vulnerability to global warming. Nature Ecology & Evolution, 4(3), Article 3. https://doi.org/10.1038/s41559-020-1114-9

Michaelides, S., Cornish, N., Griffiths, R., Groombridge, J., Zajac, N., Walters, G. J., Aubret, F., While, G. M., & Uller, T. (2015). Phylogeography and Conservation Genetics of the Common Wall Lizard, Podarcis muralis, on Islands at Its Northern Range. Plos One, 10(2), Article 2. https://doi.org/10.1371/journal.pone.0117113

Michaelides, S., While, G. M., Bell, C., & Uller, T. (2013). Human introductions create opportunities for intra-specific hybridization in an alien lizard. Biological Invasions, 15(5), Article 5. https://doi.org/10.1007/s10530-012-0353-3

Miller, R., & Plessow, F. (2013). Transformation techniques for cross-sectional and longitudinal endocrine data: Application to salivary cortisol concentrations. Psychoneuroendocrinology, 38(6), 941–946. https://doi.org/10.1016/j.psyneuen.2012.09.013

Moore, M. P., Whiteman, H. H., & Martin, R. A. (2019). A mother’s legacy: The strength of maternal effects in animal populations. Ecology Letters, 22(10), 1620–1628. https://doi.org/10.1111/ele.13351

Mueller, C. A., Eme, J., Manzon, R. G., Somers, C. M., Boreham, D. R., & Wilson, J. Y. (2015). Embryonic critical windows: Changes in incubation temperature alter survival, hatchling phenotype, and cost of development in lake whitefish (Coregonus clupeaformis). Journal of Comparative Physiology B, 185(3), 315–331. https://doi.org/10.1007/s00360-015-0886-8

Niklas, K. J., & Hammond, S. T. (2014). Assessing Scaling Relationships: Uses, Abuses, and Alternatives. International Journal of Plant Sciences, 175(7), 754–763. https://doi.org/10.1086/677238

Noble, D. W. A. (2021). MetabR/MR.R at master · daniel1noble/metabR. GitHub. https://github.com/daniel1noble/metabR

Nord, A., & Nilsson, J.-Å. (2011). Incubation temperature affects growth and energy metabolism in blue tit nestlings. The American Naturalist, 178(5), Article 5.

Osland, M. J., & Feher, L. C. (2020). Winter climate change and the poleward range expansion of a tropical invasive tree (Brazilian pepper—Schinus terebinthifolius). Global Change Biology, 26(2), 607–615. https://doi.org/10.1111/gcb.14842

Packard, G. C., Packard, M. J., Miller, K., & Boardman, T. J. (1988). Effects of temperature and moisture during incubation on carcass composition of hatchling snapping turtles (Chelydra serpentina). Journal of Comparative Physiology B, 158(1), 117–125. https://doi.org/10.1007/BF00692735

Pettersen, A. K. (2020). Countergradient variation in reptiles: Thermal sensitivity of developmental and metabolic rates across locally adapted populations. Frontiers in Physiology. https://doi.org/10.32942/osf.io/t9u8z

Pettersen, A. K., White, C. R., Bryson-Richardson, R. J., & Marshall, D. J. (2018). Does the cost of development scale allometrically with offspring size? Functional Ecology, 32(3), Article 3. https://doi.org/10.1111/1365-2435.13015

Pettersen, A. K., White, C. R., Bryson-Richardson, R. J., & Marshall, D. J. (2019). Linking life-history theory and metabolic theory explains the offspring size-temperature relationship. Ecology Letters, 22(3), Article 3. https://doi.org/10.1111/ele.13213

R Core Team. (2020). R: A language and environment for statistical computing. R Foundation for Statistical Computing, Vienna, Austria. https://www.R-project.org/.

Radder, R. S., Warner, D. A., Cuervo, J. J., & Shine, R. (2007). The Functional Significance of Residual Yolk in Hatchling Lizards Amphibolurus muricatus (Agamidae). Functional Ecology, 21(2), 302–309.

Rodríguez-Díaz, T., & Braña, F. (2012). Altitudinal variation in egg retention and rates of embryonic development in oviparous Zootoca vivipara fits predictions from the cold-climate model on the evolution of viviparity. Journal of Evolutionary Biology, 25(9), Article 9. https://doi.org/10.1111/j.1420-9101.2012.02575.x

Ruiz Miñano, M. (2021). Causes and Consequences of Hybridization – From Behaviour to Evolution. University of Tasmania.

Ruiz Miñano, M., While, G. M., Yang, W., Burridge, C. P., Salvi, D., & Uller, T. (2022). Population genetic differentiation and genomic signatures of adaptation to climate in an abundant lizard. Heredity, 128(4), 271–278. https://doi.org/10.1038/s41437-022-00518-0

Ruuskanen, S., & Hsu, B.-Y. (2018). Maternal Thyroid Hormones: An Unexplored Mechanism Underlying Maternal Effects in an Ecological Framework. Physiological and Biochemical Zoology, 91(3), Article 3. https://doi.org/10.1086/697380

Ruuskanen, S., Hsu, B.-Y., Heinonen, A., Vainio, M., Darras, V. M., Sarraude, T., & Rokka, A. (2018). A new method for measuring thyroid hormones using nano-LC-MS/MS. Journal of Chromatography B, 1093–1094, 24–30. https://doi.org/10.1016/j.jchromb.2018.06.052

Sarraude, T., Hsu, B.-Y., Groothuis, T., & Ruuskanen, S. (2020). Testing the short-and long-term effects of elevated prenatal exposure to different forms of thyroid hormones. PeerJ, 8, e10175. https://doi.org/10.7717/peerj.10175

Schulte, U., Hochkirch, A., Lötters, S., Rödder, D., Schweiger, S., Weimann, T., & Veith, M. (2012). Cryptic niche conservatism among evolutionary lineages of an invasive lizard. Global Ecology and Biogeography, 21(2), 198–211. https://doi.org/10.1111/j.1466-8238.2011.00665.x

Shine, R. (1989). Ecological influences on the evolution of vertebrate viviparity. In D. B. Wake & G. Roth (Eds.), Complex organismal functions: Integration and evolution in vertebrates (pp. 263–278). Wiley.

Shine, R. (1995). A new hypothesis for the evolution of viviparity in reptiles. The American Naturalist, 145(5), Article 5.

Shine, R. (1999). Egg-laying reptiles in cold climates: Determinants and consequences of nest temperatures in montane lizards. Journal of Evolutionary Biology, 12(5), 918–926. https://doi.org/10.1046/j.1420-9101.1999.00093.x

Sindaco, R., Razzetti, E., Doria, G., & Bernini, F. (2006). Atlas of Italian amphibians and reptiles. Edizioni Polistampa Firenze.

Sinervo, B., & Huey, R. (1990). Allometric Engineering: An Experimental Test of the Causes of Interpopulational Differences in Performance. Science (New York, N.Y.), 248, 1106–1109. https://doi.org/10.1126/science.248.4959.1106

Suarez, M. E., Wilson, H. R., Mcpherson, B. N., Mather, F. B., & Wilcox, C. J. (1996). Low Temperature Effects on Embryonic Development and Hatch Time1. Poultry Science, 75(7), 924–932. https://doi.org/10.3382/ps.0750924

Valcu, C.-M., Scheltema, R. A., Schweiggert, R. M., Valcu, M., Teltscher, K., Walther, D. M., Carle, R., & Kempenaers, B. (2019). Life history shapes variation in egg composition in the blue tit Cyanistes caeruleus. Communications Biology, 2(1), 1–14. https://doi.org/10.1038/s42003-018-0247-8

Van Damme, R., Bauwens, D., Braña, F., & Verheyen, R. F. (1992). Incubation Temperature Differentially Affects Hatching Time, Egg Survival, and Hatchling Performance in the Lizard Podarcis muralis. Herpetologica, 48(2), Article 2.

While, G. M., Williamson, J., Prescott, G., Horvathova, T., Fresnillo, B., Beeton, N. J., Halliwell, B., Michaelides, S., & Uller, T. (2015). Adaptive responses to cool climate promotes persistence of a non-native lizard. Proceedings of the Royal Society B-Biological Sciences, 282(1803), Article 1803. https://doi.org/10.1098/rspb.2014.2638

Williams, C. M., Szejner-Sigal, A., Morgan, T. J., Edison, A. S., Allison, D. B., & Hahn, D. A. (2016). Adaptation to Low Temperature Exposure Increases Metabolic Rates Independently of Growth Rates. Integrative and Comparative Biology, 56(1), Article 1. https://doi.org/10.1093/icb/icw009

Yampolsky, L. Y., & Scheiner, S. M. (1996). Why Larger Offspring at Lower Temperatures? A Demographic Approach. The American Naturalist, 147(1), 86–100.

