## Supplementary information for "Population divergence in maternal investment and embryo energy use and allocation reveals adaptive responses to cool climates"

**Table S1.** Soil temperature data for source populations of low-, mid- and high- altitude regions in Italy for *Podarcis muralis*. Data sourced for soil bioclimatic layers (0-5cm and 5-15cm depth) from Google Earth Engine global maps of soil temperature (Lembrechts et al., 2022).

| | Low | Mid | High |
|---|---|---|---|
| Mean latitude/longitude (decimal degrees) | 42.070°/ 11.954° | 42.153°/ 12.860° | 42.279°/ 13.189° |
| Mean elevation (m) | 133 | 607 | 977 |
| Annual mean temperature (°C) | | | |
| 0-5cm | 14.69 | 11.07 | 7.93 |
| 5-15cm | 14.44 | 11.73 | 9.52 |
| Mean diurnal range (°C) | | | |
| 0-5cm | 1.39 | 3.76 | 4.29 |
| 5-15cm | 4.86 | 7.17 | 6.40 |
| Max. temperature of warmest month (°C) | | | |
| 0-5cm | 22.98 | 19.52 | 18.66 |
| 5-15cm | 24.90 | 23.47 | 22.92 |
| Min. temperature of coldest month (°C) | | | |
| 0-5cm | 8.87 | 3.37 | -0.31 |
| 5-15cm | 5.55 | 2.20 | 0.07 |

**Table S2.** Sample sizes for responses to incubation temperature for embryos across different source populations (Region), Incubation treatment, and clutches (clutch order, $1^{st}$ to $5^{th}$). Number of mothers for which response data was collected shown in parentheses.

| Response | Region | Incubation treatment (°C) | Sample size |
|---|---|---|---|
| 1a. Clutch number | | | |
| | High | - | $1^{st}$ = 13, $2^{nd}$ = 11, $3^{rd}$ = 9, $4^{th}$ = 5, $5^{th}$ = 0 |
| | Mid | - | $1^{st}$ = 31, $2^{nd}$ = 29, $3^{rd}$ = 24, $4^{th}$ = 17, $5^{th}$ = 7 |
| | Low | - | $1^{st}$ = 13, $2^{nd}$ = 14, $3^{rd}$ = 14, $4^{th}$ = 12, $5^{th}$ = 4 |
| 1b. Clutch size, 1c. Clutch mass | | | |
| | High | - | 38 (14) |
| | Mid | - | 108 (32) |
| | Low | - | 57 (15) |
| 2a. Egg mass | | | |
| | High | - | 34 (14) |
| | Mid | - | 78 (32) |
| | Low | - | 39 (15) |
| 2b. Embryo retention | | | |
| | High | - | 19 (12) |
| | Mid | - | 29 (29) |
| | Low | - | 19 (13) |
| 2c. Thyroid (T4) hormone concentration, 2d. Thyroid (T3) hormone concentration, 2e. Development time | | | |
| | High | - | 29 (12) |
| | Mid | - | 68 (29) |
| | Low | - | 32 (13) |
| 3a. Development time | | | |
| | High | 21 | 32 (14) |
| | Mid | 21 | 80 (32) |
| | Low | 21 | 43 (15) |
| | High | 27 | 30 (14) |
| | Mid | 27 | 90 (31) |
| | Low | 27 | 48 (15) |
| 3b. Metabolic rate 1 (*MR*) | | | |
| | High | 21 | 11 (10) |
| | Mid | 21 | 34 (30) |
| | Low | 21 | 15 (12) |
| | High | 27 | 11 (10) |
| | Mid | 27 | 29 (30) |
| | Low | 27 | 15 (12) |
| 3c. Metabolic rate 2 ($MR_T$) | | | |
| | High | 21 | 22 (13) |
| | Mid | 21 | 59 (31) |
| | Low | 21 | 28 (14) |
| | High | 27 | 19 (13) |

| | | | |
|---|---|---|---|
| | Mid | 27 | 61(31) |
| | Low | 27 | 30 (14) |
| 4a. Tissue:total embryo mass | | | |
| | High | 21 | 5 (4) |
| | Mid | 21 | 13 (7) |
| | Low | 21 | 10 (8) |
| | High | 27 | 6 (6) |
| | Mid | 27 | 23 (17) |
| | Low | 27 | 9 (8) |
| 4b. Hatchling mass, 4c. Hatchling length | | | |
| | High | 21 | 33 (14) |
| | Mid | 21 | 88 (32) |
| | Low | 21 | 48 (15) |
| | High | 27 | 31 (14) |
| | Mid | 27 | 99 (31) |
| | Low | 27 | 52 (15) |

**Table S3.** Model ranking and selection using conditional Akaike Information Criterion (AICc) values for **1. Reproductive output** (a. Clutch number, b. Clutch size, c. Clutch mass), **2. Maternal investment** (a. Egg mass, b. Embryo retention, c. Thyroid (T4) hormone concentration, d. Thyroid (T3) hormone concentration), **3. Embryo energy use** (a. Development time, b. Metabolic rate 1 (single incubation temperature), c. Metabolic rate 2 (thermal reaction norm)), and **4. Embryo energy allocation** (a. Tissue:total embryo mass b. Hatchling mass, c. Hatchling length). Models for each response variable were ranked according to AICc value and ΔAICc is relative to best-fitting model. Model predictors (fixed effects) are provided in Table 1 in the main text.

| Response | Model number | df | LogLik | AICc | Δ AICc | Weight |
|---|---|---|---|---|---|---|
| 1a. Clutch number | 2 | 5 | -80.80 | 172.8 | 0.00 | 0.555 |
| | 1 | 3 | -83.67 | 173.8 | 1.02 | 0.333 |
| | 4 | 7 | -79.84 | 176.0 | 3.19 | 0.112 |
| 1b. Clutch size | 1 | 3 | -36.32 | 79.4 | 0.00 | 0.992 |
| | 3 | 7 | -35.46 | 89.2 | 9.77 | 0.007 |
| | 5 | 9 | -34.76 | 95.0 | 15.57 | 0.000 |
| 1c. Clutch mass | 1 | 3 | 8.98 | -11.1 | 0.00 | 0.972 |
| | 3 | 7 | 11.23 | -4.0 | 7.16 | 0.027 |
| | 5 | 9 | 11.47 | 2.9 | 14.03 | 0.001 |
| 2a. Egg mass | 1 | 4 | 356.99 | -705.7 | 0.00 | 1 |
| | 3 | 9 | 342.11 | -665.2 | 40.58 | 0 |
| | 5 | 11 | 334.77 | -646.0 | 59.77 | 0 |
| 2b. Embryo retention | 3 | 8 | -157.465 | 333.4 | 0.00 | 0.473 |
| | 5 | 10 | -154.849 | 333.6 | 0.21 | 0.425 |
| | 1 | 4 | -163.913 | 336.5 | 3.06 | 0.102 |
| 2c. Thyroid (T4) hormone | 5 | 10 | -185.947 | 393.8 | 0.00 | 0.698 |
| | 3 | 8 | -189.397 | 396.0 | 2.24 | 0.228 |
| | 1 | 4 | -194.971 | 398.3 | 4.51 | 0.073 |
| 2d. Thyroid (T3) hormone | 5 | 10 | -327.774 | 677.4 | 0.00 | 0.593 |
| | 3 | 8 | -330.481 | 678.2 | 0.75 | 0.407 |
| | 1 | 4 | -349.311 | 706.9 | 29.53 | 0.000 |
| 3a. Development time | 11 | 13 | -830.133 | 1687.4 | 0.00 | 0.974 |
| | 9 | 10 | -836.983 | 1694.7 | 7.23 | 0.026 |
| | 7 | 4 | -862.174 | 1732.5 | 45.03 | 0.000 |
| 3b. Metabolic rate (1) | 7 | 4 | 500.772 | -993.1 | 0.00 | 1 |
| | 9 | 9 | 459.952 | -900.0 | 93.18 | 0 |
| | 11 | 12 | 446.772 | -866.1 | 127.07 | 0 |
| 3c. Metabolic rate (2) | 12 | 15 | -209.576 | 449.8 | 0.00 | 1 |
| | 10 | 10 | -245.584 | 511.5 | 61.67 | 0 |
| | 8 | 4 | -274.817 | 557.7 | 107.89 | 0 |
| 4a. Tissue:total embryo mass | 9 | 9 | 77.501 | -133.7 | 0.00 | 0.623 |
| | 7 | 3 | 69.494 | -132.6 | 1.13 | 0.354 |
| | 11 | 12 | 78.540 | -127.1 | 6.65 | 0.022 |
| 4b. Hatchling mass | 9 | 8 | 72.631 | -126.5 | 0.00 | 0.811 |
| | 11 | 3 | 64.557 | -122.7 | 3.79 | 0.122 |
| | 7 | 11 | 74.441 | -121.5 | 5.00 | 0.067 |
| 4c. Hatchling length | 11 | 13 | -773.310 | 1573.8 | 0.00 | 0.802 |
| | 9 | 10 | -777.949 | 1576.6 | 2.80 | 0.198 |
| | 7 | 4 | -869.299 | 1746.7 | 172.91 | 0.000 |

**Table S4**. Summary of parameter estimates for best-fitting linear models and linear mixed effects models (with Maternal ID as a random effect), describing the relationship between 1. Reproductive output, 2. Maternal investment, 3. Embryo energy use, and 4. Embryo energy allocation, with predictor variables: Maternal body mass, Temperature, Incubation treatment, Region, Egg mass, and Clutch number, and their interactions as shown in Table 1. Final model parameters were included based on model selection (AICc) shown in Table S2. Estimates for Region and Clutch no. are relative to 'High' altitude and '2nd' clutch factor levels, respectively. Significant estimates shown in bold: $*p < 0.05$, $**p < 0.01$, $***p < 0.001$.

| | Estimate | SE | t | df | *p* |
|---|---|---|---|---|---|
| **1a. Clutch number (m2)** | | | | | |
| Intercept | 3.29 | 0.90 | 3.65 | 1 | **<0.001*** ** |
| Maternal body mass | -0.09 | 0.21 | -0.43 | 1 | 0.67 |
| Region | Low: 1.12 | 0.47 | 2.37 | 2 | **0.02*** |
| | Mid: 0.74 | 0.43 | 1.72 | | 0.09 |
| **1b. Clutch size (m1)** | | | | | |
| Intercept | 2.86 | 0.73 | 3.95 | 1 | **<0.001*** ** |
| Maternal body mass | 0.34 | 0.14 | 2.45 | 1 | **0.02*** |
| **1c. Clutch mass (m1)** | | | | | |
| Intercept | 0.50 | 0.19 | 2.61 | 1 | **0.01*** |
| Maternal body mass | 0.17 | 0.04 | 4.54 | 1 | **<0.001*** ** |
| **2a. Egg mass (m1)** | | | | | |
| Intercept | 0.22 | 0.02 | 12.54 | 1 | **<0.001*** ** |
| Maternal body mass | 0.01 | 0.00 | 4.21 | 1 | **<0.001*** ** |
| **2b. Embryo retention (m3)** | | | | | |
| Intercept | 25.26 | 1.81 | 13.11 | 1 | **<0.001*** ** |
| Maternal body mass | 0.37 | -0.36 | 0.86 | 1 | 0.39 |
| Region | Low: -2.31 | 1.09 | -2.12 | 2 | **0.04*** |
| | Mid: -1.21 | 0.95 | -1.27 | | 0.22 |
| Clutch no. | 3rd: -1.24 | 0.76 | -1.65 | 2 | 0.11 |
| | 4th: -0.90 | 0.89 | -1.02 | | 0.31 |
| **2c. [Thyroid hormone (T4)] (m5)** | | | | | |
| Intercept | 3.52 | 1.22 | 2.88 | 1 | **<0.01*** ** |
| Maternal body mass | 0.15 | 0.29 | 0.50 | 1 | 0.62 |
| Region | Low: 6.72 | 2.80 | 2.40 | 2 | **0.02*** |
| | Mid: 0.05 | 1.42 | 0.04 | | 0.97 |
| Clutch no. | 3rd: 0.49 | 0.20 | 2.49 | 2 | **0.01*** |
| | 4th: 0.80 | 0.24 | 3.35 | | **<0.01*** ** |
| Maternal body mass × Region | Low: -1.26 | 0.55 | -2.29 | 2 | **0.02*** |
| | Mid: 0.05 | 0.33 | 0.14 | | 0.89 |
| **2d. [Thyroid hormone (T3)] (m5)** | | | | | |
| Intercept | 5.80 | 4.00 | 1.45 | 1 | 0.15 |
| Maternal body mass | 0.56 | 0.96 | 0.58 | 1 | 0.56 |
| Region | Low: 6.73 | 9.16 | 0.73 | 2 | 0.46 |
| | Mid: 0.53 | 4.65 | 0.11 | | 0.91 |
| Clutch no. | 3rd: 2.90 | 0.64 | 4.56 | 2 | **<0.001*** ** |
| | 4th: 3.73 | 0.76 | 4.90 | | **<0.001*** ** |
| Maternal body mass × Region | Low: -1.23 | 1.80 | -0.68 | 2 | 0.50 |
| | Mid: 0.10 | 1.07 | 0.09 | | 0.93 |
| **2e. Development time (m6; 21 °C)** | | | | | |
| Intercept | 90.91 | 6.50 | 13.99 | 1 | **<0.001*** ** |
| [T4] | -0.07 | 0.61 | -0.11 | 1 | 0.91 |
| [T3] | -0.12 | 0.22 | -0.54 | 1 | 0.59 |
| Stage at oviposition | -0.24 | 0.21 | -1.16 | 1 | 0.25 |

| | | | | | |
|---|---|---|---|---|---|
| **2e. Development time (m6; 27 °C)** | | | | | |
| Intercept | 44.59 | 1.80 | 24.74 | 1 | **<0.001*** ** |
| [T4] | 0.00 | 0.18 | 0.01 | 1 | 1.00 |
| [T3] | 0.00 | 0.06 | 0.04 | 1 | 0.97 |
| Stage at oviposition | -0.21 | 0.06 | -3.37 | 1 | **<0.01** ** |
| **3a. Development time (m11)** | | | | | |
| Intercept | 88.62 | 2.76 | 32.14 | 1 | **<0.001*** ** |
| Incubation treatment | 27 °C: -48.51 | 3.68 | -13.17 | 1 | **<0.001*** ** |
| Region | Low: 4.48 | 1.00 | 4.64 | 2 | **<0.001*** ** |
| | Mid: 1.66 | 0.90 | 1.84 | | 0.07 |
| Egg mass | -23.35 | 9.67 | -2.41 | 1 | **0.02*** |
| Clutch number | 3$^{rd}$: -0.91 | 0.54 | -1.71 | 2 | 0.09 |
| | 4$^{th}$: -1.03 | 0.55 | -1.88 | | 0.06 |
| Incubation treatment × Region | 27 °C × Low: -2.31 | 1.36 | -1.69 | 1 | 0.09 |
| | 27 °C × Mid: -0.54 | 1.23 | -0.44 | | 0.66 |
| Incubation treatment × Egg mass | 27 °C: 18.82 | 12.71 | 1.48 | 1 | 0.14 |
| **3b. Metabolic rate (MR; m7)** | | | | | |
| Intercept | $8.78\times10^{-3}$ | $2.50\times10^{-4}$ | 35.08 | 1 | **<0.001*** ** |
| Incubation treatment | 27 °C: $5.11\times10^{-3}$ | $3.63\times10^{-4}$ | 14.09 | 1 | **<0.001*** ** |
| **3c. Metabolic rate ($MR_T$; m12)** | | | | | |
| Intercept | -6.10 | 0.28 | -21.86 | 1 | **<0.001*** ** |
| Incubation treatment | 27 °C: 0.86 | 0.42 | 2.05 | 1 | **0.04*** |
| Region | Low: 0.04 | 0.08 | 0.52 | 2 | 0.60 |
| | Mid: -0.10 | 0.07 | -1.42 | | 0.16 |
| Egg mass | 0.64 | 0.19 | 3.33 | 1 | **<0.001*** ** |
| Temperature | 0.12 | 0.00 | 52.37 | 1 | **<0.001*** ** |
| Clutch number | 2$^{nd}$: -0.50 | 0.11 | -4.48 | 3 | **<0.001*** ** |
| | 3$^{rd}$: -0.40 | 0.11 | -3.36 | | **<0.001*** ** |
| | 4$^{th}$: -0.23 | 0.12 | -1.92 | | 0.06 |
| Incubation treatment × Region | 27 °C × Low: -0.25 | 0.03 | -1.95 | 2 | 0.05 |
| | 27 °C × Mid: 0.01 | 0.11 | 0.14 | | 0.89 |
| Incubation treatment × Egg mass | 27 °C: -0.07 | 0.31 | -0.22 | 1 | 0.83 |
| Incubation treatment × Temperature | 27 °C: -0.03 | 0.00 | -7.61 | 1 | **<0.001*** ** |
| **4a. Tissue:total embryo mass (m9)** | | | | | |
| Intercept | 0.60 | 0.10 | 6.18 | 1 | **<0.001*** ** |
| Incubation treatment | 27 °C: -0.06 | 0.02 | -2.79 | 1 | **<0.01** ** |
| Region | Mid: 0.08 | 0.02 | 3.40 | 2 | **<0.01** ** |
| | High: 0.09 | 0.03 | 2.71 | | **<0.01** ** |
| Egg mass | -0.57 | 0.33 | -1.72 | 1 | 0.09 |
| Clutch number | 3$^{rd}$: 0.03 | 0.02 | 1.18 | 2 | 0.24 |
| | 4$^{th}$: 0.00 | 0.03 | 0.11 | | 0.91 |
| **4b. Hatchling mass (m9)** | | | | | |
| Intercept | 0.13 | 0.02 | 7.72 | 1 | **<0.001*** ** |
| Incubation treatment | 27 °C: -0.01 | 0.00 | -2.99 | 1 | **<0.01** ** |
| Region | Low: 0.02 | 0.01 | 2.20 | 2 | **0.03*** |
| | Mid: 0.01 | 0.01 | 0.96 | | 0.34 |
| Egg mass | 0.82 | 0.06 | 14.63 | 1 | **<0.001*** ** |
| Clutch number | 2$^{nd}$: -0.02 | 0.00 | -3.92 | 3 | 0.46 |
| | 3$^{rd}$: -0.00 | 0.00 | -0.74 | | 0.62 |
| | 4$^{th}$: -0.00 | 0.01 | -0.49 | | 0.92 |
| **4c. Hatchling length (m11)** | | | | | |
| Intercept | 47.30 | 2.40 | 19.70 | 1 | **<0.001*** ** |
| Incubation treatment | 27 °C: 4.41 | 2.87 | 1.54 | 1 | 0.13 |
| Region | Low: 5.15 | 1.07 | 4.82 | 2 | **<0.001*** ** |

| | | | | | |
|---|---|---|---|---|---|
| | Mid: 2.21 | 0.93 | 2.38 | | **0.02*** |
| Egg mass | 51.03 | 8.47 | 6.02 | 1 | **<0.001***** |
| | $2^{nd}$: -0.41 | 0.46 | -0.90 | | 0.37 |
| Clutch number | $3^{rd}$: -0.33 | 0.53 | -0.65 | 3 | 0.52 |
| | $4^{th}$: 0.26 | 0.55 | 0.49 | | 0.62 |
| Incubation treatment × Region | 27 °C × Low: 1.01 | 1.08 | 0.93 | 2 | 0.35 |
| | 27 °C × Mid: 1.56 | 0.95 | 1.21 | | 0.23 |
| Incubation treatment × Egg mass | 27 °C: -6.07 | 10.11 | -0.60 | 1 | 0.55 |

**Table S5**. Parameter estimates (± standard error; SE) from nonlinear regression of metabolic rate (*MR*) and temperature (T) across three altitude regions (High, Mid, Low) and incubation treatments (21 °C and 27 °C), where $MR = f \times \exp^{(a \times T)}$. Significance level *$p < 0.05$, **$p < 0.01$, ***$p < 0.001$.

| Region | Incubation treatment (°C) | Estimate intercept (*f*) | SE | t-value | *p* | Estimate slope (*a*) | SE | t-value | *p* |
|---|---|---|---|---|---|---|---|---|---|
| High | 21 | $4.827 \times 10^{-3}$ | $7.661 \times 10^{-4}$ | 6.30 | <0.0001*** | 0.649 | 0.058 | 11.22 | <0.0001*** |
| | 27 | $7.953 \times 10^{-3}$ | $1.358 \times 10^{-3}$ | 5.85 | <0.0001*** | 0.538 | 0.063 | 8.492 | <0.0001*** |
| Mid | 21 | $4.453 \times 10^{-3}$ | $5.492 \times 10^{-4}$ | 8.11 | <0.0001*** | 0.668 | 0.045 | 14.92 | <0.0001*** |
| | 27 | $7.536 \times 10^{-3}$ | $8.139 \times 10^{-4}$ | 9.26 | <0.0001*** | 0.532 | 0.042 | 13.24 | <0.0001*** |
| Low | 21 | $6.148 \times 10^{-3}$ | $1.233 \times 10^{-3}$ | 4.99 | <0.0001*** | 0.585 | 0.074 | 7.909 | <0.0001*** |
| | 27 | $6.316 \times 10^{-3}$ | $1.108 \times 10^{-3}$ | 5.70 | <0.0001*** | 0.552 | 0.065 | 8.503 | <0.0001*** |

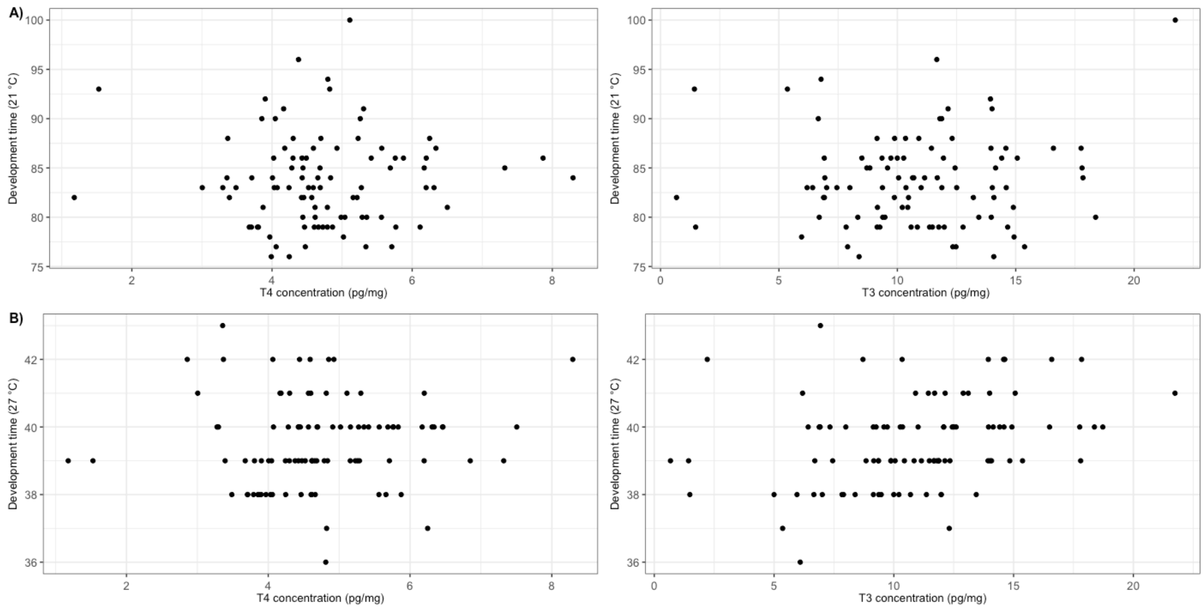


**Figure S1.** Relationship between yolk thyroid hormone (T4 and T3) concentration and development time (from oviposition until hatching) at A) 21°C and B) 27 °C. Data points represent raw data. No significant correlation was found after accounting for stage at oviposition.
